# Altered follicular immunity in secondary lymphoid organs is associated with interferon hyperactivity in Down syndrome

**DOI:** 10.64898/2026.08.04.741562

**Authors:** Jeremías Dutto, Jeremias Bustos, Lucia Boffelli, Jimena Tosello Boari, Jenny Christine Kienzler, Paula Araya, Sabrina Dhooge, Ana Flores Guirado, Pilar Biasi, Ruth Eliana Baigorri, Carina Valeriani, Wilfrid Richer, Cecilia del Carmen Montes, Virginia Cecconi, Burkhard Becher, Joaquín M. Espinosa, Eliane Piaggio, Nicolás Gonzalo Nuñez, Mariana Maccioni

## Abstract

Down syndrome, caused by trisomy 21, is characterized by chronic interferon-associated inflammation and immune dysregulation, yet the contribution of human secondary lymphoid organs to shaping this immune landscape remains unclear. Using multimodal single-cell and spatial profiling of human tonsils, we identify extensive remodeling of immune organization in trisomy 21. CD4⁺ T cells are skewed away from canonical follicular helper (TFH) programs toward inflammatory TFH1-like and cytotoxic helper states enriched for interferon-responsive transcriptional programs. Tonsillar TFH cells exhibit increased interferon-γ and interleukin-21 production, indicating inflammatory skewing toward type 1 helper immunity. These alterations are accompanied by changes in dendritic cell and CD8⁺ T-cell compartments, reduced follicular size, increased extrafollicular TFH1–B-cell proximity and altered B-cell differentiation trajectories. Together, our findings identify trisomy 21 as a unique human context to investigate how chronic interferon-associated inflammation reshapes lymphoid tissue organization and adaptive immune cell fate decisions.

## Main

Down syndrome (DS), caused by trisomy 21 (T21), is associated with developmental delay and a broad spectrum of co-occurring conditions, including increased susceptibility to infections, autoimmunity and leukemia, alongside a reduced incidence of solid tumors ^1–3^. These clinical manifestations reflect profound immune dysregulation affecting both innate and adaptive immunity from early life ^4–6^. However, the molecular mechanisms underlying the altered disease susceptibility in DS remain incompletely understood ^7,8^.

Accumulating evidence indicates that many of these immune abnormalities are linked to triplication of four interferon receptor genes (*IFNAR1, IFNAR2, IFNGR2* and *IL10RB*) encoded on chromosome 21, resulting in increased receptor expression, heightened sensitivity to interferon signaling and chronic interferon-associated inflammation ^9–12^. Indeed, individuals with DS exhibit persistent activation of interferon-stimulated genes and systemic inflammatory signatures that have led to the proposal that DS presents similarities to genetically determined human interferonopathies ^9,11,12^.

Current knowledge of immune dysfunction in DS is largely derived from studies performed in peripheral blood^4,5^. These investigations have demonstrated hyperresponsive CD4⁺ T cells, resistance to regulatory T-cell mediated suppression, skewing toward TH1/TH17 differentiation programs, and features of activation and senescence within CD8⁺ T-cell compartment^4,5,13–15^. Similarly, B-cell immunity is intrinsically perturbed, with reduced class-switched memory, impaired somatic hypermutation and defects in humoral responses ^5,14,16–18^. However, whether these peripheral immune abnormalities originate from, or are reinforced by, altered immune organization within human secondary lymphoid organs (SLOs)—the site where adaptive immune responses are initiated, coordinated and refined—remains largely unknown.

SLOs are highly organized microenvironments in which interactions among dendritic cells (DCs), T cells and B cells coordinate adaptive immune responses, germinal center (GC) reactions and long-term immune memory^19,20^. Increasing evidence indicates that chronic inflammatory environments can profoundly remodel cellular interactions within lymphoid tissues, altering immune cell positioning, differentiation trajectories and humoral immunity^19,21^. In general, these phenomena have been described in chronic viral infections and autoimmune diseases such as systemic lupus erythematosus, where sustained interferon signaling is associated with disruption of canonical GC responses and expansion of inflammatory and extrafollicular pathways ^21–28^ Yet, how chronic interferon-biased inflammation affects the organization and function of human SLOs remains poorly understood.

Given the central role of CD4⁺ T cells in orchestrating adaptive immunity, particularly through the generation and maintenance of germinal center responses, we hypothesized that chronic interferon-driven inflammation in DS remodels immune cell interactions within SLOs, altering CD4⁺ T-cell differentiation and follicular immunity and ultimately contributing to the adaptive immune abnormalities observed in the circulation. To address this question, we focused on tonsils, a strategically positioned SLO continuously exposed to environmental antigens and highly enriched in germinal centers^29^. We combined single-cell transcriptomic, proteomic and immune receptor profiling with multiplex immunofluorescence analyses of non-infectious tonsils from control children (D21) and children with T21.

Here, we show that chronic interferon-driven inflammation in T21 is associated with profound remodeling of CD4⁺ T-cell differentiation within human tonsils, characterized by expansion of inflammatory TFH1-like and cytotoxic helper states together with reduced canonical follicular programs. These alterations are accompanied by changes in DCs composition, CD8⁺ T-cell states, B-cell differentiation trajectories and reduced follicular organization, suggesting extensive rewiring of cellular interactions within human SLO.

T21 therefore represents a genetically defined human setting of sustained interferon-associated inflammation, offering unique insights into how persistent inflammatory cues reshape immune cell differentiation and tissue organization. More broadly, these principles may extend beyond DS and be relevant to a wide range of interferon-driven inflammatory and autoimmune diseases.

### Single-cell profiling reveals altered immune composition and dendritic cell remodeling in tonsils from children with trisomy 21

To define how T21 reshapes immune organization and CD4⁺ T-cell differentiation within SLOs, we generated a single-cell multi-omic atlas of non-infectious tonsils from control children (D21, n=3) and children with DS (T21, n=3) using CITE-seq with paired TCR/BCR sequencing. To enable robust detection of CD4⁺ T-cell heterogeneity in this B cell–rich tissue, we partially reduced B-cell predominance by mixing sorted CD19⁺ and CD19⁻ fractions at a 30:70 ratio, while preserving representation of major immune compartments (see Methods; **Fig. 1A**). Key observations were then validated by flow cytometry (D21, n=8; T21, n=15) and mIF. Multi-modal computational approaches were subsequently applied to integrate transcriptomic, proteomic and immune repertoire information (**Fig. 1A**).

**Figure 1:**
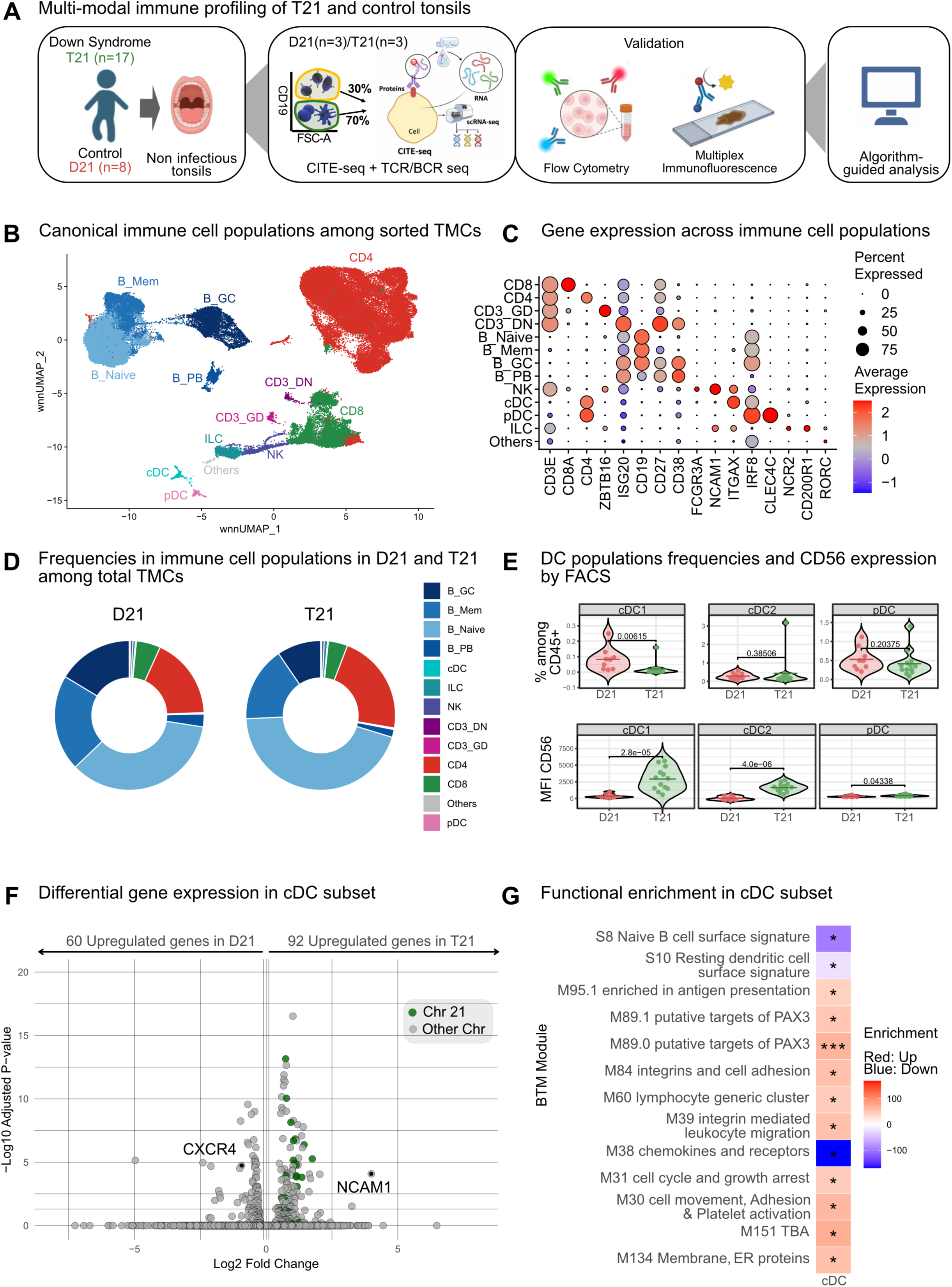
Single-cell profiling reveals altered immune composition and dendritic cell remodeling in tonsils from children with trisomy 21. **(A)** Study design and analytical workflow. Non-infectious tonsils from individuals with trisomy 21 (T21) and euploid controls (D21) were subjected to multi-modal immune profiling. Discovery analyses were performed by CITE-seq coupled to paired TCR/BCR sequencing in sorted tonsillar mononuclear cells (TMCs) from D21 (n = 3) and T21 (n = 3) individuals. To enrich for T cells and improve the representation of minor lymphocyte populations, CD19⁺ and CD19⁻ fractions were initially sorted and subsequently recombined at a 30:70 ratio prior to CITE-seq analysis. Findings were further validated in an expanded cohort by flow cytometry (D21, n = 8; T21, n = 15) and multiplex immunofluorescence **(B)** Weighted nearest-neighbor (wnn) UMAP embedding of CD45⁺ TMCs (92,350 cells) integrating transcriptomic and surface protein information after mixing CD19^-^ and CD19^+^ cells in a 30:70 ratio. Major immune populations are annotated, including CD4⁺ and CD8⁺ T cells, B-cell subsets: naïve (B_Naïve), memory (B_Mem), germinal center (B_GC) and plasma cells (B_PC), NK cells, innate lymphoid cells (ILC), conventional dendritic cells (cDC), plasmacytoid dendritic cells (pDC) and minor immune populations. **(C)** Dot plot showing representative marker genes used for cluster annotation across immune populations. Dot size indicates the percentage of cells expressing each gene, and color intensity indicates scaled average expression. **(D)** Relative frequencies of major immune populations determined by flow cytometry in unfractionated TMCs prior to CD19-based enrichment and CITE-seq analysis (D21, n=3 ; T21 n=3). **(E)** Flow cytometry quantification of different DC subsets in independent tonsil samples from the whole cohort mentioned in (A). Each dot represents a sample (upper panel). Mean Fluorescence Intensity (MFI) of NCAM⁺ (CD56) in the different DC populations (lower panel). P values were calculated using the Mann–Whitney–Wilcoxon test. **(F)** Volcano plot showing differential gene expression between cDC cells (cDC1 and cDC2) from tonsils of children with T21 and D21, highlighting the expression levels of NCAM1 and CXCR4. Green dots represent chromosome 21 encoded genes (Log2FC>0.2, and adj.pVal<0.05). **(G)** Heatmap showing significant enrichment of Blood Transcriptional Modules (BTMs) across cDC cells (cDC1 and cDC2) comparing T21 and D21 samples. Color intensity represents the direction and magnitude of enrichment (red: upregulated in T21; blue: downregulated in T21). Significant modules were defined using p < 0.05 corrected for multiple testing by false discovery rate q < 0.05 according to Li et al., 2014 ^32^. *\*\*\*p<0.001; **p<0.01; *p<0.05*

We first defined global changes in immune composition and antigen-presenting compartments that may shape CD4⁺ T cell differentiation. Weighted nearest neighbour (wnn) integration resolved major immune populations, integrating ∼21,000 transcripts and 137 surface proteins across 92,350 high-quality tonsillar cells (**Fig. 1B-C, Extended Data Fig. 1A**). Following the recently published tonsil atlas^29^ and based on gene and protein expression, we annotated 13 canonical immune populations, including CD8⁺ and CD4⁺, γδ, CD4^-^CD8^-^ (CD3_DN) T cells, naïve (B_Naïve), memory (B_Mem), germinal center B (B_GC) cells, and (B_PCB plasmablasts, NK cells, conventional DCs (cDCs), plasmacytoid DCs (pDCs) as well as innate lymphoid cells (ILCs) and other minor populations (**Fig. 1B-C and Extended Data Fig. 1A**). To determine global immune composition in T21 tonsils, we first quantified major immune populations in unfractionated tonsillar mononuclear cells (TMCs) by flow cytometry prior to cell sorting. This analysis revealed an overall preservation of major immune lineages between D21 and T21 samples, with only modest shifts in the relative abundance of B- and T-cell compartments (**Fig. 1D and Extended Data Fig. 1B**). Consistent with these observations, flow cytometric validation in the expanded cohort showed a tendency toward increased T-cell frequencies together with a reduction in B cells (x ± SD, B cells: D21 71±12 vs T21 80±9, p=0.076; T cells: D21 19±8 vs T21 28±12, p=0.076) whereas the CD4/CD8 ratio remained largely preserved **(Extended Data Fig. 1B-C)**.

To characterize the innate immune landscape and identify innate populations potentially involved in shaping CD4⁺ T cell differentiation in T21 tonsils, we subclustered innate-lineage cells and visualized them using wnn-UMAP (**Extended Data Fig. 1D-E**). This analysis identified discrete clusters corresponding to NK cells, ILCs, CD3_DN T cells, CD3_GD T cells, cDCs, and pDCs. The global structure of the innate compartment was largely conserved between D21 and T21 samples. However, the density of cells across clusters suggested compositional shifts within NK and ILC populations in T21 tonsils (**Extended Data Fig. 1D**). To achieve higher resolution despite the limited sample size, we applied Milo analysis to test for differential abundance across local neighbourhoods of phenotypically similar cells. This approach enables the detection of subtle population shifts that may not be captured by conventional clustering strategies. Neighbourhood-based differential abundance testing identified a small number of differentially abundant NK- and ILC-associated neighbourhoods in T21 tonsils **(Extended Data Fig. 1F),** indicating subtle changes within these compartments. Similarly, no major differences were detected among cDC or pDC subsets, likely reflecting the limited number of myeloid cells captured by the single-cell dataset. Given the central role of DCs cells in instructing CD4⁺ T-cell differentiation ^30,31^, we next examined DC subsets by flow cytometry in the expanded cohort (Extended Data Fig. 1G), enabling more sensitive detection of differences in these relatively rare populations. (**Extended Data Fig. 1G**). Flow cytometry analyses revealed selective remodeling of DC subsets within the CD45⁺ compartment (**Fig. 1E).** The frequency of cDC1 (CD141⁺CD1c⁻) cells was significantly reduced among CD45+ cells in T21, whereas cDC2 (CD141⁻CD1c⁺) and pDC frequencies were preserved. Notably, CD56 expression (*NCAM*) was elevated across all DC subsets from T21 patients (**Fig. 1E, lower panel**). Transcriptomic profiling of combined cDCs (cDC1 and cDC2) revealed a distinct molecular signature in T21 tonsils, with upregulation of several chromosome 21–encoded genes consistent with dosage effects, along with transcriptional changes in genes beyond chromosome 21 (**Fig. 1F**). *NCAM* upregulation at the mRNA level mirrored protein data, supporting increased CD56⁺ expression in DC subsets. Conversely, *CXCR4* expression was severely downregulated. To define the immune pathways underlying the observed transcriptional changes, we performed enrichment analysis using blood transcriptional modules (BTMs)^32^. BTMs are co-expression–based gene sets derived from blood transcriptomes that summarize coordinated immune pathways and facilitate systems-level analysis of gene expression data. BTM analysis further indicated functional reprogramming of cDCs from T21 tonsils (**Fig. 1G**). Reduced expression of the S10 resting DC program together with enrichment of the M95.1 antigen-presentation module indicated a transition toward an activated DC state. In parallel, attenuation of the M38 migration/adhesion signature, largely attributable to diminished *CXCR4* expression, suggested disrupted positioning dynamics within SLOs.

Collectively, these data reveal that tonsils from T21 individuals exhibit a remodelled DC landscape marked by cDC1 reduction, increased CD56 expression across DC subsets, and diminished *CXCR4* expression, features predicted to influence CD4⁺ T cell priming and differentiation within SLOs^31,33,34^. Given the well-established role of DC in integrating inflammatory and interferon signals to instruct TFH differentiation, these findings raised the possibility that the altered DC landscape in T21 tonsils may contribute to the rewiring of downstream CD4⁺ T-cell trajectories.

### Trisomy 21 promotes interferon-associated memory, cytotoxic and helper programs at the expense of canonical TFH differentiation

To test this hypothesis, we next investigated the impact of T21 on tonsillar CD4⁺ T-cell differentiation within SLOs. Differential expression analysis revealed that T21 CD4⁺ T cells upregulate TH1-associated genes, including *IFNG, STAT4, IRF1, TBX21* and *CXCR3*, consistent with a pro-inflammatory, interferon-biased program (**Fig. 2A**). BTM analysis further demonstrated enrichment of modules related to type I interferon response pathways (M127, M75), T cell activation (M7.0, M7.3), MAPK signalling (M100), cytokine responses (M24), supporting an interferon-biased inflammatory program (**Fig. 2B**).

**Figure 2:**
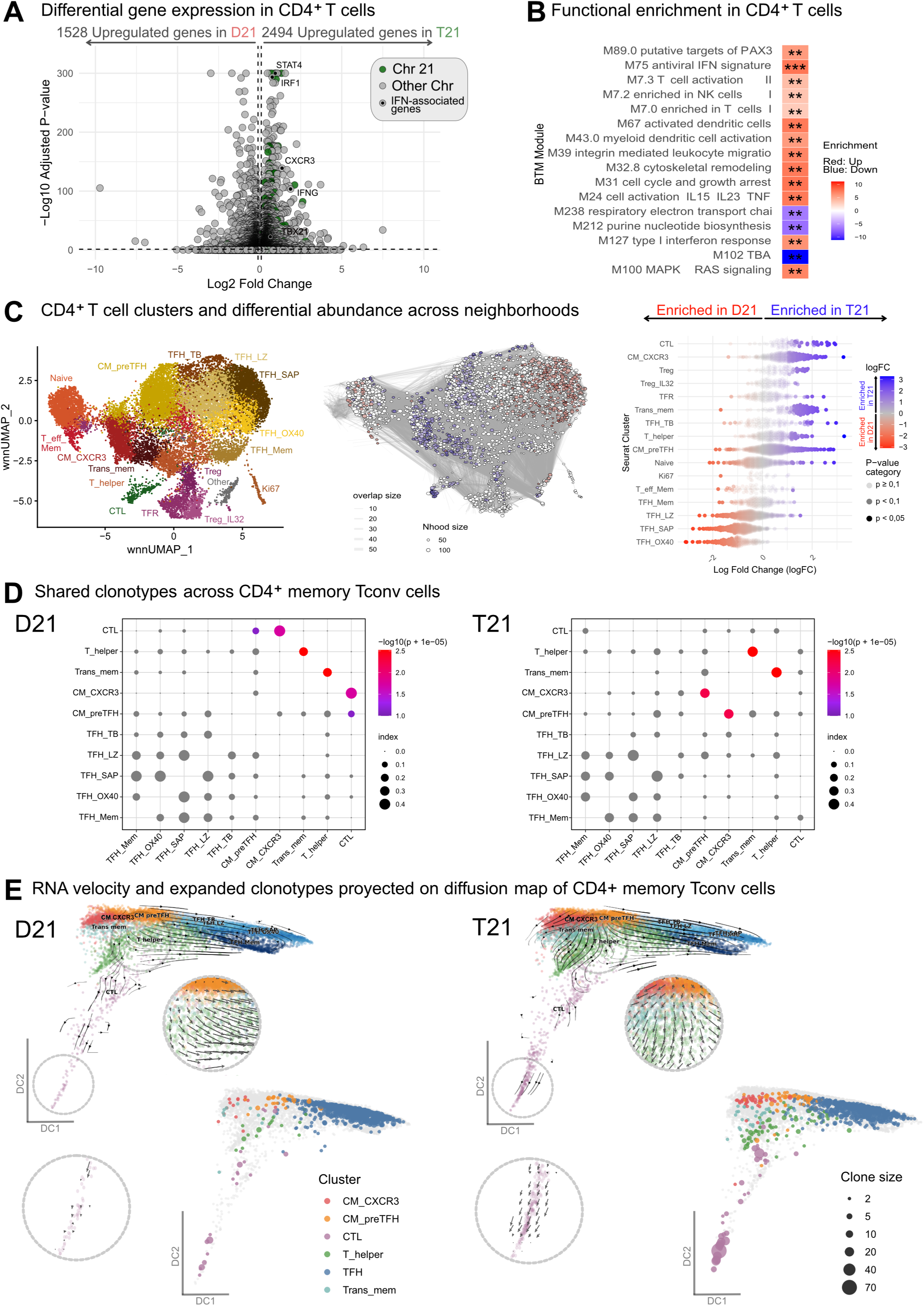
Trisomy 21 reshapes tonsillar CD4⁺ T-cell differentiation toward interferon-associated memory programs at the expense of canonical TFH states. **(A)** Volcano plot showing differentially expressed genes between tonsillar CD4⁺ T cells from children with T21 and D21. Green dots indicate chromosome 21-encoded genes and black dots represent interferon-associated genes. Selected genes associated with TH1 polarization and inflammatory responses, including *STAT4*, *IRF1*, *CXCR3*, *IFNG* and *TBX21*, are highlighted. **(B)** Heatmap showing significant enrichment of BTMs across CD4⁺ T cells comparing T21 and D21 samples. The heatmap shows modules significantly enriched in the T21 versus D21 as described in Fig. 1G. *\*\*\*p<0.001; **p<0.01* **(C)** wnnUMAP representation of tonsillar CD4⁺ T cells colored according to transcriptionally defined clusters, including naïve, central memory CXCR3⁺ (CM_CXCR3), CM_preTFH, transitional memory (Trans_mem), T_helper, effector memory (T_eff_Mem), cytotoxic (CTL), proliferating (Ki67), Treg, Treg_IL32, TFR and multiple TFH states (TFH_Mem, TFH_LZ, TFH_TB, TFH_OX40 and TFH_SAP). **Middle panel:** Milo neighborhood graph, where each node represents a local transcriptional neighborhood and edge connectivity reflects transcriptional similarity between neighborhoods. Node color indicates differential abundance between conditions (logFC), with blue representing enrichment in T21 and red enrichment in D21. **Right panel:** Beeswarm plot with the distribution of log fold changes (logFC) in neighborhood abundance identified by Milo across annotated CD4^+^ T cell clusters. Each point represents a neighborhood assigned to a cluster, and the x-axis shows the log fold change in abundance between T21 and D21 samples. Point color indicates the magnitude and direction of the change, while point size reflects statistical significance . **(D)** TCR clonotype sharing analysis across T conv (non-naïve, non-Treg CD4⁺ T-cell clusters) in D21 and T21 tonsils. Dot plots represent pairwise Morisita-Horn similarity indices based on shared *TRB* clonotypes. Dot size indicates the degree of clonotype overlap between clusters, while color intensity denotes statistical significance. **(E)** RNA velocity analysis of tonsillar non naïve,-non Treg CD4⁺ T-cell states in D21 and T21 samples using diffusion maps and wnnUMAP embeddings. Black streamlines indicate the inferred direction of transcriptional transitions. Insets highlight differences in local differentiation trajectories between conditions. Lower panels show expanded clonotypes projected onto diffusion maps, with dot size proportional to clone size.

Projection of tonsillar CD4+ T cells onto the reference atlas from Massoni-Badosa *et al*^29^ resolved 16 transcriptionally distinct populations spanning naïve, effector memory (T_eff_Mem), central memory expressing *CXCR3* (CM_CXCR3), transitional memory (Trans-mem, an intermediate state between CM and fully differentiated T-Eff-Mem cells), T_helper, CM_preTFH, multiple TFH states (TFH-TB; TFH-LZ; TFH-SAP, TFH-OX40, TFH-Mem), regulatory populations (Treg, Treg_ IL32 and TFR), and proliferating cells (*MKI67*) (**Fig. 2C left panel and Extended data Fig. 2A and 2B**). Among these, we identified a rare ZEB2⁺GZMH⁺CD4⁺ T cell population (CTL) expressing *CXCR3* and *IFNG*, consistent with a terminally differentiated cytotoxic helper-like state previously associated with chronic interferon exposure and autoimmune inflammation^35–37^. Milo analysis revealed extensive remodeling of CD4⁺ T cell neighbourhoods in T21 tonsils, with enrichment of CTL, CM_CXCR3+, CM_pre TFH, Trans-mem, T helper, and TFH-TB subsets and some neighbourhoods from Tregs, alongside relative contraction of canonical GC-associated TFH populations (TFH-LZ, TFH-SAP, and TFH-OX40) (**Fig. 2C middle and right panel**).

To investigate lineage relationships underlying the remodeling of conventional memory CD4⁺ T cells in T21 tonsils, we combined clonal overlap and RNA velocity analyses across transcriptionally defined states. Conventional memory CD4⁺ T cells (memory Tconv) were defined as non-naïve, non-Treg CD4⁺ T cells. Morisita-based repertoire analysis revealed extensive clonotype sharing among memory, helper and TFH-associated populations in both D21 and T21 tonsils, with CM_preTFH cells representing a major hub of clonal connectivity linking memory and effector populations (CM_CXCR3, Trans_Mem, T_helper and CTL) with follicular states (TFH_mem, TFH_OX40, TFH_SAP and TFH_LZ) (**Fig. 2D).** However, the architecture of clonal relationships differed markedly between conditions. In D21 tonsils, CM_preTFH cells displayed broad repertoire overlap with Trans_memory, helper and TFH populations, functioning as a central node connecting inflammatory and follicular compartments. In contrast, T21 tonsils showed preferential clonotype sharing between CM_preTFH and the inflammatory CM_CXCR3 population, accompanied by markedly reduced repertoire overlap with TFH subsets.

Consistent with these clonal relationships, RNA velocity projected a continuous differentiation landscape connecting CM_CXCR3 and CM_preTFH populations with downstream TFH-associated states in both conditions (**Fig. 2E**). Velocity confidence scores were uniformly high across the major differentiation trajectories, supporting robust velocity estimates throughout the manifold **(Extended Data Fig. 2C).** Whereas D21 trajectories were largely organized around transitions toward follicular states, T21 cells displayed a broader differentiation landscape spanning helper, memory and follicular compartments, suggesting increased differentiation plasticity (**Fig. 2E, regions highlighted by circles**). Consistent with these findings, UpSet analyses demonstrated a substantial increase in the size of the CM_CXCR3 and CM_preTFH clonotype sets in T21 tonsils while clonotype intersections spanning multiple TFH clusters and CM_preTFH cells are evident in both conditions **(Extended Data Fig. 2D)**. Mapping of expanded clonotypes onto diffusion maps (**Fig. 2E, lower panel)** further revealed that clonally expanded cells in T21 occupied broader regions of the differentiation landscape and preferentially localized to CM_CXCR3⁺, CM_preTFH and CTL populations, whereas expanded clones in D21 appeared more restricted to follicular-associated states. Together, these findings support a model in which T21 promotes the diversification and expansion of clonally related CD4⁺ T cells toward inflammatory, cytotoxic and T_helper trajectories at the expense of canonical follicular programs.

### Functional validation of interferon-biased CD4⁺ T-cell remodeling and emergence of TFH1-like states in trisomy 21 tonsils

To determine whether the CD4⁺ T-cell remodeling identified by CITE-seq could be detected in the whole cohort, we quantified phenotypically related Tconv CD4⁺ populations by flow cytometry (**Fig. 3A and Extended Data Fig. 3A)**. Consistent with the overall trends observed by single-cell analyses, T21 tonsils displayed a significant reduction in naïve and TFH CD4⁺ T cells together with an expansion of CXCR3⁺ memory populations. On the other hand, Treg frequencies remained unchanged (**Fig. 3A**). To investigate whether this transcriptional remodeling translated into functional alterations, we quantified ex vivo cytokine production by CD4⁺ T-cell subsets. Non-follicular CD4⁺ T cells exhibited only minor changes in inflammatory molecule production between conditions (**Extended Data Fig. 3B**). In contrast, TFH cells from T21 tonsils displayed a selective increase in IL21⁺IFNγ⁺ double-producing cells, consistent with the emergence of TFH1-like functional states (**Fig. 3B)**. These findings indicate that the interferon-associated programs identified by single-cell analyses are accompanied by functional polarization of follicular helper cells.

**Figure 3:**
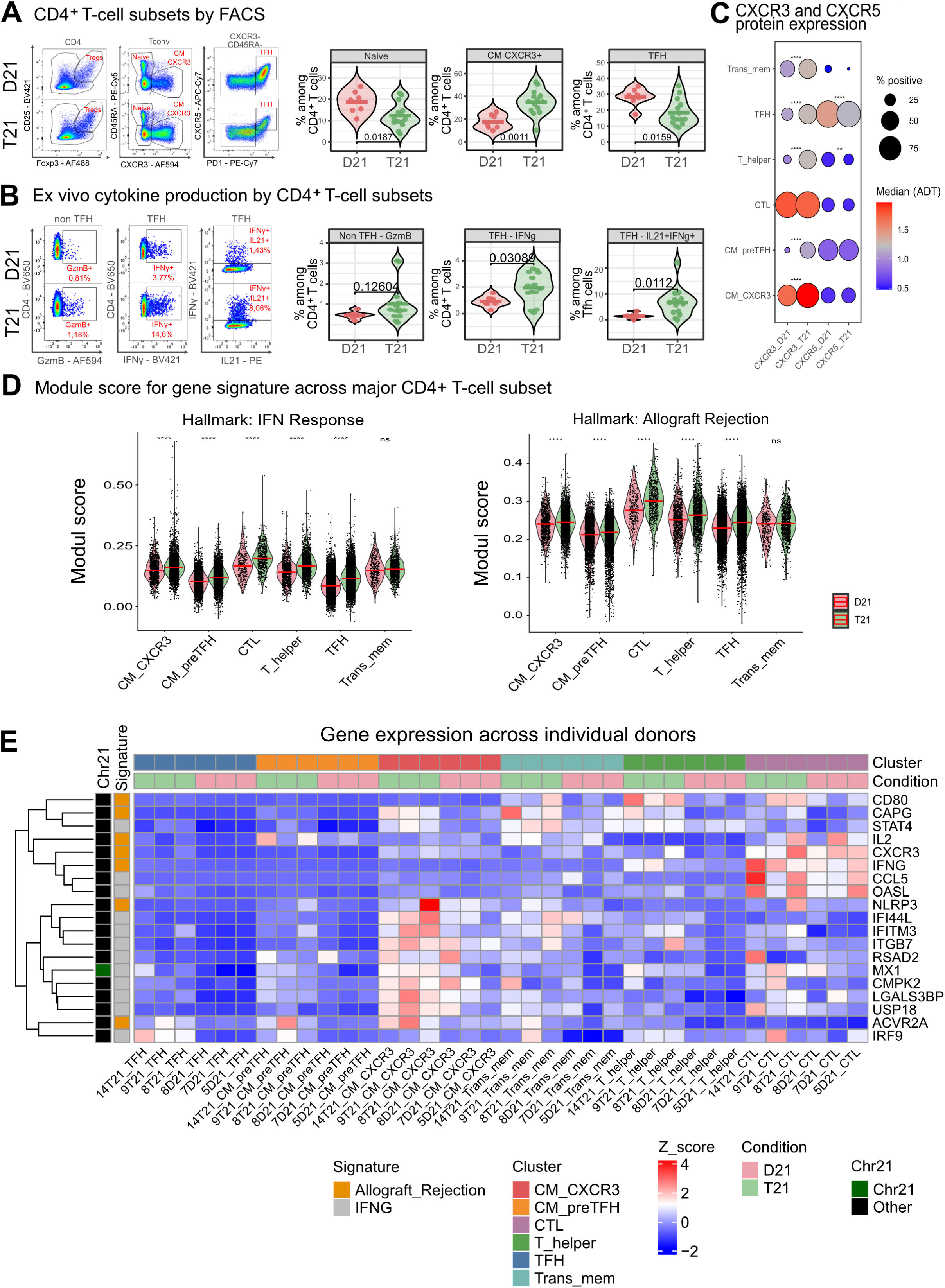
Functional validation of interferon-biased CD4⁺ T-cell remodeling and emergence of TFH1-like states in trisomy 21 tonsils. A) Flow cytometric validation of major conventional CD4⁺ T-cell populations in the whole cohort of tonsils (D21, n = 8; T21, n = 15). Representative gating strategy showing identification of regulatory T cells (Tregs; CD25⁺FoxP3⁺) and conventional CD4⁺ T cells (Tconv; CD25⁻FoxP3⁻), followed by delineation of naïve (CD45RA⁺), CXCR3⁺ memory-like (CM_CXCR3⁺), and TFH (PD1⁺CXCR5⁺) populations within the Tconv compartment. Violin plots depict frequencies of the indicated populations among total CD4⁺ T cells. P values were calculated using two-tailed Mann–Whitney tests. **(B)** Functional characterization of CD4⁺ T-cell cytokine production in tonsils. Representative flow cytometry plots and quantification of granzyme B production in non-TFH cells and IFNγ production in TFH cells are shown. Frequencies of IL21⁺IFNγ⁺ double-producing TFH cells are depicted on the right. Each dot represents one individual. P values were calculated using two-tailed Mann–Whitney tests. **C)** CITE-seq-derived protein expression of CXCR3 and CXCR5 across major CD4⁺ T-cell populations. Dot size represents the percentage of positive cells and color intensity indicates median ADT expression. **D)** Module scores for Hallmark IFNγ response and Allograft Rejection gene signatures across major CD4⁺ T-cell populations. Individual cells are shown as dots and distributions are summarized by violin plots. P values were calculated using the Mann–Whitney– Wilcoxon test. *\*\*\* P<0.001; ***** P<0.0001, ns= non significant* **E) H**eatmap depicting scaled pseudobulk expression of top differential expressed genes from the Hallmark IFNγ response and Allograft Rejection signatures across individual donors and major conventional CD4⁺ T-cell subsets. Columns correspond to donor-subset combinations and rows to individual genes. Top annotations indicate cell subset, donor condition (D21 or T21), and chromosome 21 gene localization. Expression values are shown as Z scores.

CITE-seq-derived protein measurements further revealed increased CXCR3 expression across multiple CD4⁺ T-cell subsets including Trans_Mem, CM_CXCR3, CM_preTFH, T_helper, CTL and TFH cells, together with reduced CXCR5 expression, most prominently within the TFH compartment (**Fig. 3C and Extended Data Fig. 3C).** Together, these observations support a shift away from canonical follicular programs toward interferon-associated helper states. To define population-specific transcriptional alterations, we performed differential expression analyses within individual CD4⁺ T-cell clusters **(Extended Data Fig. 3D)**. TFH, CM_preTFH, T_helper, CTL, Trans_mem, and CM_CXCR3 subsets consistently upregulated interferon-responsive genes, including *IFI44L*, *IFITM3*, *IRF1*, *STAT4* and *CXCR3*, together with chromosome 21-encoded genes such as *RUNX1* and *MX1*. Notably, the TFH compartment showed coordinated upregulation of interferon-responsive genes including *IFNG, STAT4*, *IRF1* and *CXCR3*, supporting the enrichment of TFH1-like states in T21 tonsils. This transcriptional profile was further supported by the increased frequency of IL21⁺IFNγ⁺ TFH cells detected ex vivo. In contrast, CTL cells preferentially upregulated cytotoxic effector molecules, including *CCL5*, *NKG7*, *GZMH, GZMA* and *PRF1*, consistent with their lineage identity **(Extended Data Fig. 3D)**.

Finally, the Hallmark Interferon-γ Response and Allograft Rejection gene sets were enriched across CM_CXCR3, TFH, CM_preTFH, CTL, T_helper and Trans_Mem populations from T21 tonsils, consistent with coordinated activation of interferon-responsive, antigen-presentation and cytotoxic effector programs (**Fig. 3D)**, indicating that interferon-associated inflammatory programs permeate multiple helper states rather than being restricted to a single CD4⁺ T-cell population.

To assess whether these inflammatory programs were consistently represented across individuals, we examined the expression of representative genes from the Hallmark IFNγ response and Allograft Rejection signatures in each patient and CD4⁺ T-cell cluster (**Fig. 3E)**. This analysis revealed a consistent enrichment of interferon-associated genes across T21 samples, particularly within CM_CXCR3, CM_preTFH, CTL,T_helper, TFH and Trans_mem subsets. Increased expression of canonical interferon-responsive genes, including **IFI44L, IFITM3, RSAD2, MX1, CMPK2, USP18 and IRF9**, together with TH1-associated molecules such as **STAT4, CXCR3, IFNG and CCL5**, was observed across multiple T21 donors. In contrast, TFH cells displayed comparatively weaker induction of these signatures. Thus, the IFNγ-and allograft rejection-associated transcriptional programs identified at the population level are reproducibly detected across individual T21 patients rather than being driven by outlier samples.

These findings indicate that the inflammatory transcriptional program observed in T21 is not restricted to a single CD4⁺ T-cell subset but instead permeates multiple helper and memory populations, supporting the emergence of a broadly interferon-biased immune environment in T21 tonsils.

### Trisomy 21 reshapes the tonsillar CD8⁺ T-cell compartment through expansion of inflammatory resident memory-like programs and depletion of cytokine-enriched states

Given the extensive interferon-associated remodeling observed in DCs and CD4⁺ T cells, and because CD8⁺ T cells can both respond to and reinforce inflammatory cues within SLO tissues through cytokine production and interactions with antigen-presenting cells^38^ we next investigated whether the CD8⁺ compartment was similarly remodeled in T21 tonsils. Differential expression analysis revealed increased expression of cytotoxic effector genes (*GZMB, GZMH, PRF1* and *CTSW*), inflammatory mediators (*TNF*) and *STAT4* in CD8⁺ T cells from T21 tonsils (**Fig. 4A)**. This program emerged in the absence of increased *TBX21* expression and was accompanied by reduced *EOMES* and *TOX* expression, arguing against classical exhaustion or stable terminal differentiation states. Instead, these findings suggest the acquisition of an activated inflammatory program by tonsillar CD8⁺ T cells.

**Figure 4:**
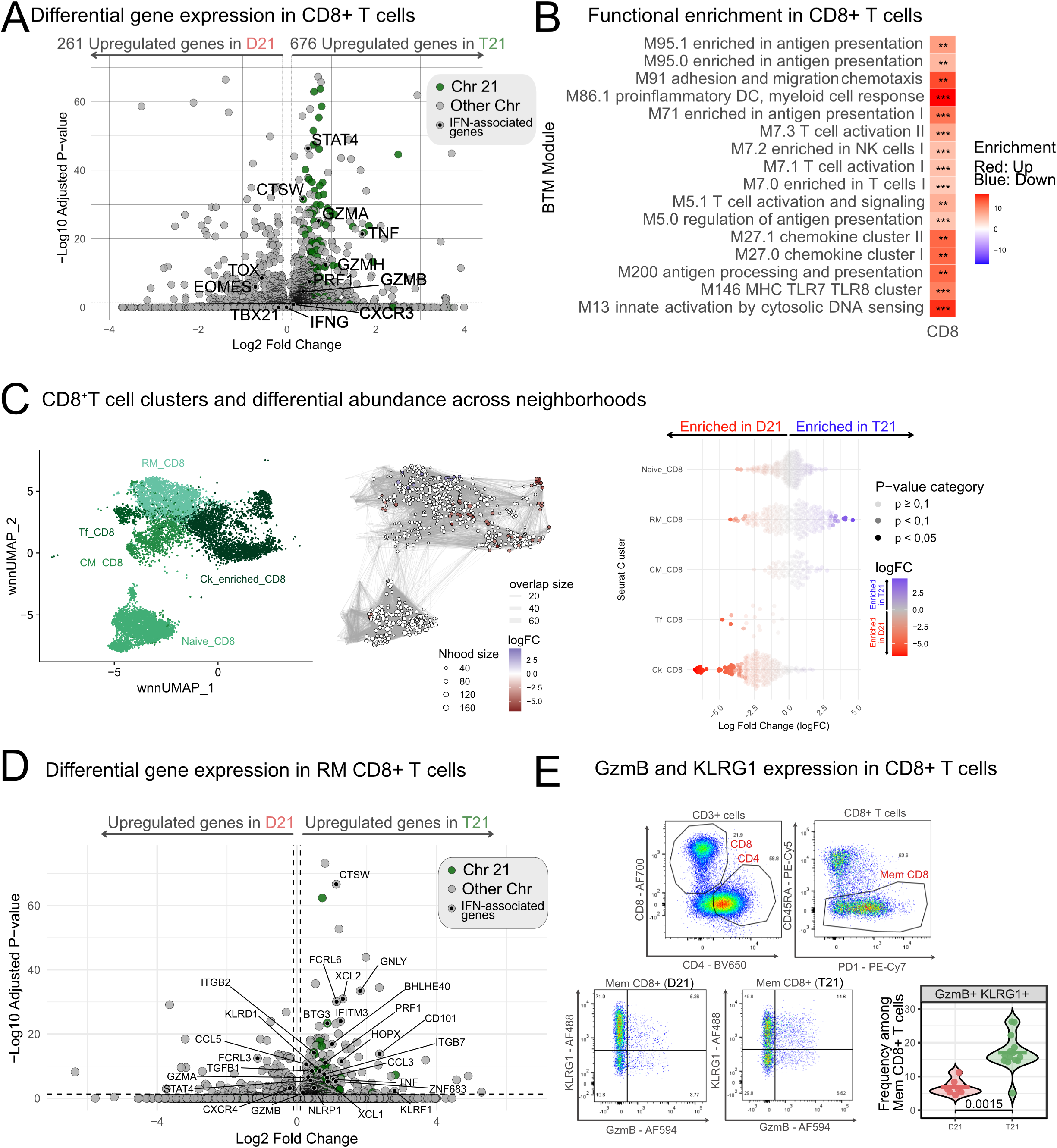
Trisomy 21 reshapes the tonsillar CD8⁺ T-cell compartment through expansion of inflammatory resident memory-like programs and depletion of cytokine-enriched states. **(A)** Volcano plot showing differentially expressed genes between total CD8⁺ T cells from D21 and T21 tonsils. Green dots represent chromosome 21-encoded genes and black dots indicate interferon-associated genes. Selected inflammatory and cytotoxic genes, including S*TAT4, CTSW, GZMA, TNF, GZMH, PRF1, IFNG and GZMB,* are highlighted. **(B)** Heatmap showing significant enrichment of BTM across CD8⁺ T cells comparing T21 and D21 tonsils. Modules associated with antigen presentation, innate activation, T-cell activation, chemokine signaling, and NK-cell programs were significantly enriched in T21-derived CD8⁺ T cells. Red indicates positive enrichment and blue negative enrichment. The heatmap shows modules significantly enriched in the T21 versus D21 as described in Fig. 1G. *\*\*\*p<0.001; **p<0.01* **C)** wnnUMAP embedding of CD8⁺ T cells from D21 and T21 tonsils integrating transcriptomic and surface protein information. CD8⁺ T cells segregate into five major populations: naïve CD8⁺ T cells (Naive_CD8), central memory CD8⁺ T cells (CM_CD8), resident memory CD8⁺ T cells (RM_CD8), follicular CD8⁺ T cells (Tf_CD8) and a chemokine-enriched effector population (Ck_CD8).**Middle panel:** Milo differential abundance analysis of CD8⁺ T cell subsets in tonsils from children with T21 and D21. **Right panel:** Beeswarm plot showing the distribution of log fold changes (logFC) in neighborhood abundance identified by Milo across annotated CD8^+^ T cell clusters. Each point represents a neighborhood from the single-cell graph assigned to a specific CD8^+^ T cell subset, including Naive_CD8, RM_CD8, CM_CD8, Tf_CD8 and Ck_CD8 T cells. Point color indicates the direction and magnitude of the logFC (red: decreased in T21; blue: increased in T21). Point color indicates the magnitude and direction of the change, while point size reflects statistical significance. **(D)** Differential gene expression analysis of RM_CD8 populations comparing T21 and D21 tonsils. T21 RM_CD8 cells upregulated genes associated with cytotoxicity, tissue residency and inflammatory activation, including *CTSW, GNLY, FCRL6, IFITM3, BHLHE40, HOPX, PRF1, GZMB, KLRF1, XCL1* and *TNF*. **(E)** Flow cytometric validation of memory CD8⁺ T cells. Representative gating strategy identifying memory CD8⁺ T cells and expression of granzyme B and KLRG1. Violin plots show increased frequencies of GzmB⁺KLRG1⁺ cells among memory CD8⁺ T cells in T21 tonsils.Each dot represents one individual. P values were calculated using two-tailed Mann–Whitney tests.

BTM enrichment analysis further demonstrated overrepresentation of antigen presentation and DC-associated inflammatory modules (M95.0, M95.1 and M86.1), largely driven by increased HLA-DR expression, together with enrichment of T-cell activation (M7.1, M7.3) and NK/cytotoxic modules (M7.2) (**Fig. 4B)**, supporting the presence of a broadly activated inflammatory CD8⁺ T-cell state.

Projection onto the human tonsil reference atlas ^29^ resolved five transcriptionally distinct populations, including naïve (Naive_CD8), central memory (CM_CD8), resident memory-like (RM_CD8), follicular (Tf_CD8) and cytokine-enriched (Ck_CD8) populations, the latter characterized by expression of *XCL1* and *XCL2* together with innate-like lymphocyte markers (**Fig. 4C** and **Extended Data Fig. 4A-B**). Milo analysis revealed remodeling of CD8⁺ T-cell neighbourhoods in T21 tonsils, characterized by changes in RM_CD8-associated neighbourhoods and marked depletion of Ck_CD8 neighbourhoods (**Fig. 4C, right panel)**, suggesting perturbation of specialized chemokine-producing CD8⁺ T-cell states. Interestingly, despite their reduced abundance, residual Ck_CD8 cells from T21 tonsils displayed additional transcriptional remodeling characterized by increased expression of *CCL3L3, CCL4L2, NCAM1, KIR2DS4* and *TGFB1*, together with reduced *CXCR4* expression **(Extended Data Fig. 4C)**, indicating acquisition of NK-like and chemokine-producing features.

In contrast, RM_CD8 cells, previously described as preferentially localized within the epithelium and subepithelial connective tissue surrounding tonsillar crypts ^29^, underwent marked qualitative remodeling in T21 tonsils. Differential expression analysis revealed increased expression of genes associated with cytotoxicity, tissue adaptation and inflammatory activation, including *CTSW, GNLY, CCL5, PRF1, TNF, KLRD1, HOPX* and *ZNF683* (**Fig. 4D)**. Consistent with these findings, flow cytometry analyses in the expanded cohort demonstrated increased frequencies of GzmB⁺KLRG1⁺ memory CD8⁺ T cells in T21 tonsils (**Fig. 4E)**.

Collectively, these findings indicate that T21 is associated with both quantitative and qualitative remodeling of the tonsillar CD8⁺ T-cell compartment, combining depletion and phenotypic reprogramming of CK-enriched CD8⁺ T cells with acquisition of inflammatory resident memory-like programs. These alterations are consistent with a broader interferon-biased immune environment across T21 tonsils.

### Interferon-associated B-cell activation uncouples from proliferative germinal center programs in trisomy 21

Given the contraction of canonical TFH cells together with the expansion of inflammatory CM_CXCR3⁺ and TFH1-like populations, we hypothesized that altered follicular T-cell help would be accompanied by changes in B-cell activation and differentiation.

Differential expression analysis revealed extensive transcriptional remodeling in B cells from T21 tonsils, characterized by increased expression of interferon- and activation-associated genes, including STAT1, STAT4, CXCR3, TNF, JUNB, NR4A1 and NR4A2, together with several chromosome 21-encoded transcripts (**Fig. 5A).** Consistent with these findings, BTM enrichment analysis demonstrated enrichment of type I interferon and antiviral pathways (M75, M68, M127, M150), AP-1 transcriptional networks (M20) and inflammatory modules, whereas several cell-cycle and mitotic programs (M4.5, M4.6, M4.10, M4.11, M6 and M8) were significantly depleted in T21 B cells (**Fig. 5B)**, suggesting activation in the absence of proportional expansion of proliferative programs.

**Figure 5:**
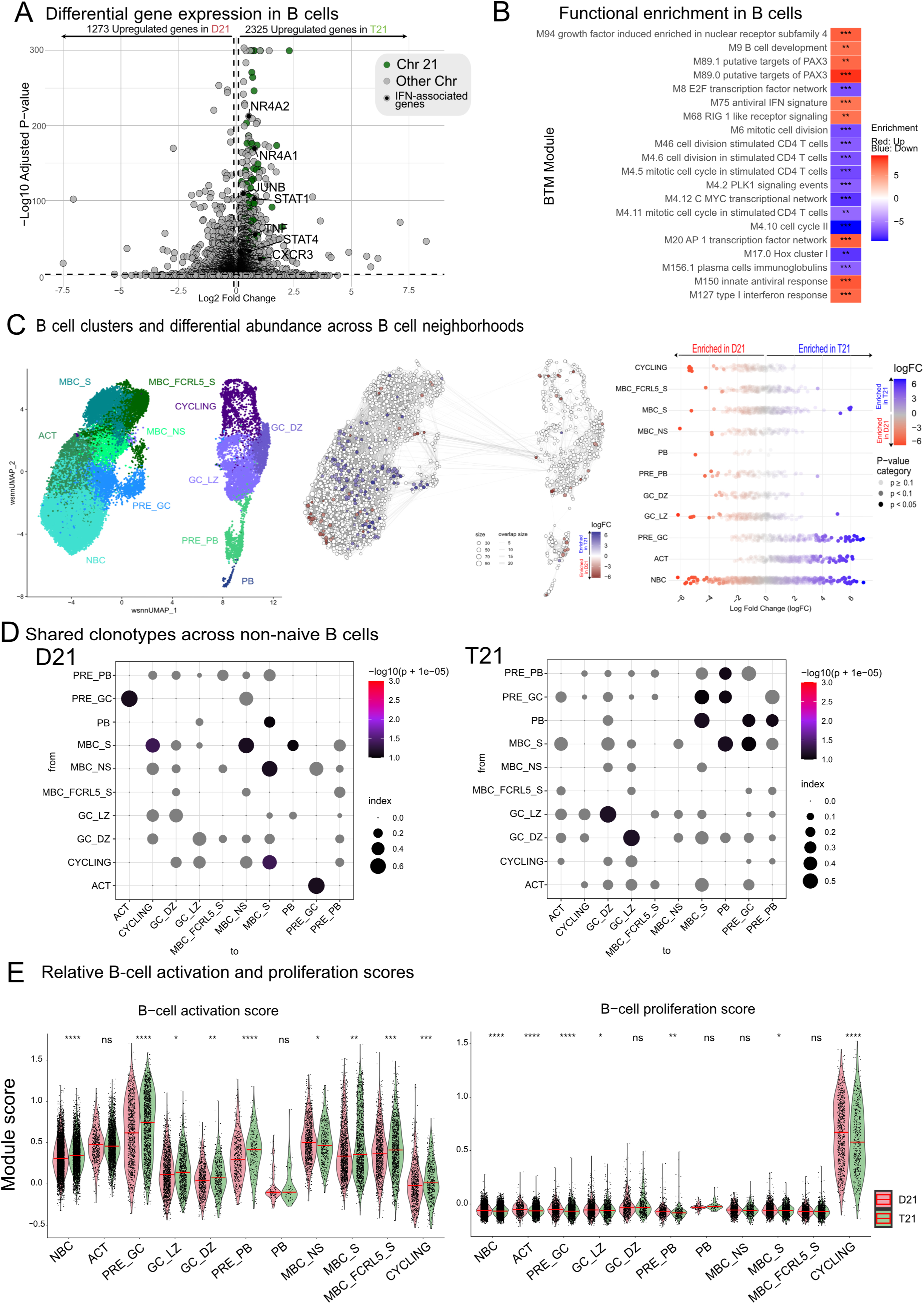
Interferon-associated B-cell activation uncouples from proliferative germinal center programs in trisomy 21. **(A)** Differential gene expression analysis of total B cells from D21 and T21 tonsils. Volcano plot showing genes differentially expressed between T21 and D21 B cells. Green dots indicate chromosome 21-encoded genes and black dots indicate interferon-associated genes. Selected genes associated with activation and inflammatory responses, including *NR4A1, NR4A2, JUNB, STAT1, TNF, STAT4* and *CXCR3*, are highlighted. **(B)** Heatmap showing significant enrichment of BTMs across total B-cells from T21 versus D21 tonsils. Red indicates positive enrichment and blue indicates negative enrichment as explained in Fig. 1G. *\*\*\*p<0.001; **p<0.01* **(C)** wnnUMAP representation of major tonsillar B-cell populations, including naïve B cells (NBC), activated B cells (ACT), pre-germinal center cells (PRE_GC), germinal center light-zone (GC_LZ) and dark-zone (GC_DZ) B cells, pre-plasmablasts (PRE_PB), plasmablasts (PB), non-switched memory B cells (MBC_NS), switched memory B cells (MBC_S), FCRL5⁺ switched memory B cells (MBC_FCRL5_S), and cycling cells. **Middle panel:** Milo neighborhood graph illustrating differential abundance between D21 and T21 tonsils. Node size is proportional to neighborhood size and node color indicates differential abundance (logFC). **Right panel:** Differential abundance analysis of B-cell populations. Beeswarm plot showing the distribution of neighborhood log fold changes across annotated B-cell populations. **(D)** Shared BCR clonotypes across B-cell clusters. Morisita-based analysis showing overlap of clonotypes among B-cell subsets in D21 and T21 tonsils. Dot size represents clonotype overlap index and color indicates significance. **(E)** Relative activation and proliferation scores across B-cell states. Module scores calculated across annotated B-cell subpopulations in D21 and T21 tonsils. Individual cells are shown as dots and distributions are summarized by violin plots.

We then projected tonsillar B cells onto the human tonsil B-cell reference atlas^29^, identifying naïve (NBC), activated (ACT), cycling, pre-germinal center (PRE_GC), germinal center light zone (GC_LZ), germinal center dark zone (GC_DZ), pre-plasmablast (PRE_PB), plasmablast (PB), switched and non-switched memory populations (MBC_S and MBC_NS), as well as FCRL5⁺ switched memory B cells (MBC_FCRL5_S), a differentiated, interferon-associated memory B-cell subset frequently expanded in chronic inflammatory settings^39^ (**Fig. 5C** and **Extended Data Fig. 5A-B)**. Milo analysis revealed selective remodeling of B-cell neighbourhood composition in T21 tonsils, characterized by enrichment of ACT and PRE_GC clusters together with relative depletion of several GC-associated neighbourhoods (**Fig. 5 C, middle and right panels)**, suggesting altered progression through canonical GC differentiation programs.

To validate these findings, we quantified major B-cell subsets by flow cytometry in the whole cohort of tonsils **(Extended Data Fig. 5C)**. Although frequencies of major B-cell subsets were largely preserved, CXCR3 expression was significantly increased across multiple B-cell subsets, including MBC_S and MBC_NS, atypical, GC and PB populations **(Extended Data Fig. 5D lower panel-E)**. These findings indicate that T21 primarily reshapes the activation state of B cells rather than causing major changes in their global composition. Also, the increased CXCR3 expression across multiple B-cell populations suggests enhanced responsiveness to inflammatory chemokine gradients and may reflect exposure to IFNγ-associated microenvironmental cues.

To determine whether altered differentiation states were accompanied by changes in clonal relationships, we next examined BCR repertoire overlap across B-cell populations. Morisita-based analysis revealed substantial clonotype sharing between ACT and PRE_GC populations in both conditions (**Fig. 5D)**. However, the overall organization of clonal relationships differed markedly between groups. In T21 tonsils, ACT subsets displayed broader clonal connectivity, with increased sharing with cycling, GC and memory subsets. Notably, the relationships between PB and PRE_PB populations also differed substantially between D21 and T21 samples. Whereas D21 PB showed clonotype sharing with GC and MBC_S populations, T21 tonsils displayed a broader PB-associated clonal network that additionally encompassed PRE_GC and PRE_PB states (**Fig. 5D)**. Consistent with these findings, UpSet analyses demonstrated a substantial increase in the size of the PRE_GC clonotype set in T21 tonsils, accompanied by higher-order intersections involving ACT, pre-GC, GC and memory states. By contrast, D21 samples displayed a more GC-centered clonal architecture **(Extended Data Fig. 5F)**.

Since BTM analysis suggested activation in the absence of proportional expansion of proliferative programs, we calculated activation and proliferation module scores across all B-cell subsets. Compared with D21 controls, T21 B cells exhibited broadly increased activation scores in most B cell subsets, particularly within PRE_GC, indicating widespread engagement of activation-associated programs (**Fig. 5E).** In striking contrast, proliferation scores were reduced across most T21 B-cell subsets, with the notable exception of GC_DZ cells, which retained proliferative features (**Fig. 5E)**. Thus, T21 B cells display a marked uncoupling between activation and proliferative programs, consistent with altered germinal center dynamics and inefficient progression through canonical GC responses.

### TFH1 polarization is associated with reduced follicular size and rewired T–B interactions in T21 tonsils

To determine whether the interferon-biased immune remodeling observed by single-cell and functional analyses was associated with altered tissue organization, we examined tonsillar architecture by multiplex immunofluorescence. T21 tonsils displayed reduced mean follicular diameter, whereas follicle density was preserved, indicating altered follicular organization rather than loss of follicular structures (**Fig. 6A)**. Across samples, follicle size positively correlated with the frequency of conventional TFH cells and negatively correlated with CM_CXCR3+ cell frequencies measured by flow cytometry, linking the balance between follicular and inflammatory helper states to tissue architecture (**Fig. 6B)**.

**Figure 6.**
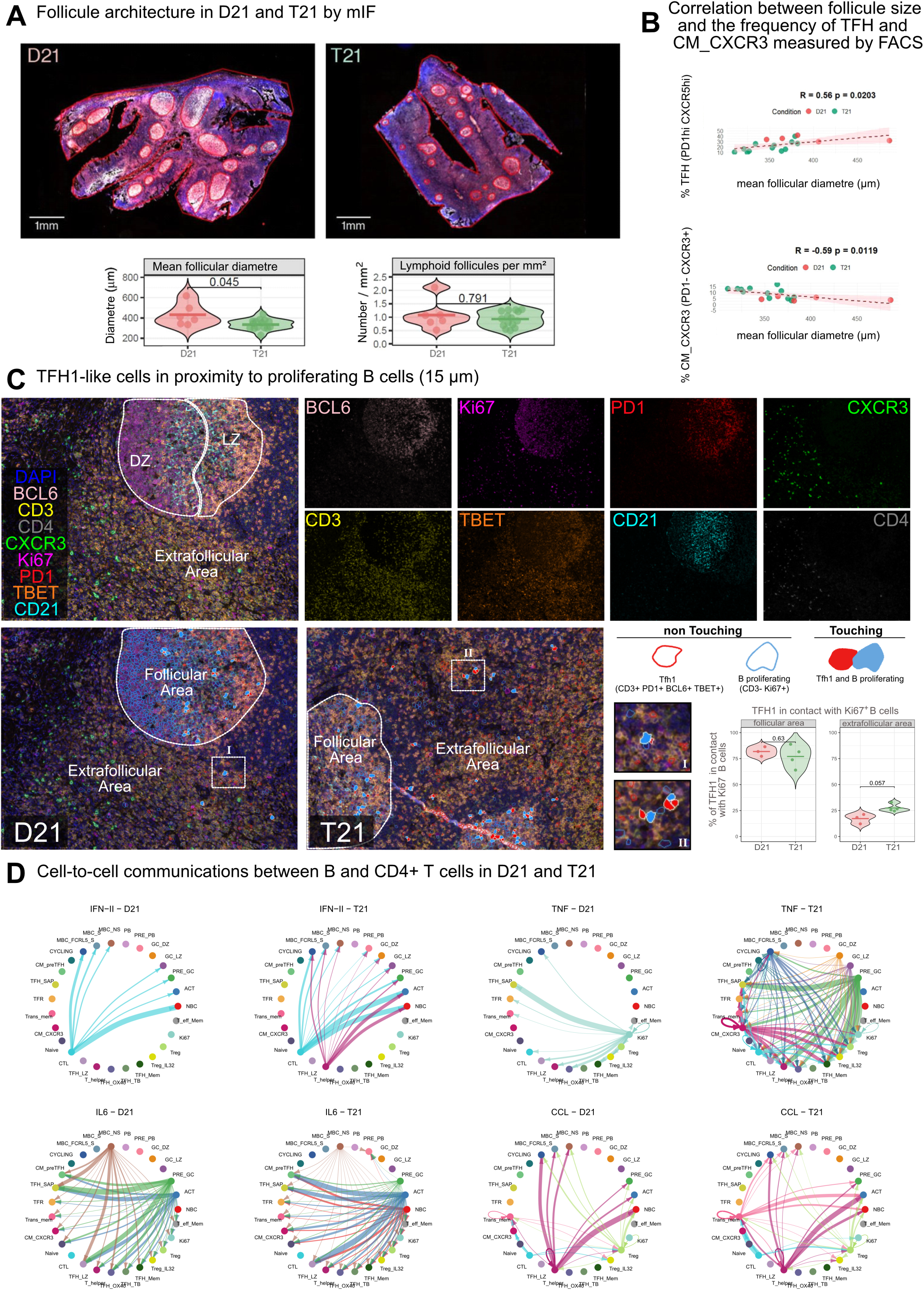
Altered follicular architecture and association with inflammatory TFH1-like states in trisomy 21 tonsils. **(A)** Representative images of multiplex immunofluorescence from FFPE tissue samples obtained from T21 (n=4) and control (n=3) tonsils. Slides stained with anti-BCL6 were scanned with PhenoImager Fusion (Akoya) and follicle size was measured with QuPath (10 to 35 follicles per slide). Each point represents the mean ± SEM of the diameter of the follicles measured in each tonsil (p<0.01) (n= 9 T21 and n= 6 D21). Statistical analysis was performed using the Unpaired t test. B) Pearson correlation analysis between the size of the follicles measured in (A) and the percentages of CM_CXCR3 and of canonical TFH cells identified by flow cytometry as shown in Fig.3A (D21, n=5; T21, n=12). C) Representative images of mIF from FFPE tissue samples obtained from T21 (n=4) and control (n=3) tonsils. The lower panel displays the various markers analyzed (BCL6, Ki67, CD3, CD4, PD1, CXCR3, and TBET). The upper panel shows merged images of these markers. Light zone (LZ) was defined by CD21 staining and dark zone (DZ) was defined by Ki67. The dotted squares (I and II) indicate the T cell zones digitally amplified to show the presence of TFH1 cells in D21 and T21 tonsils respectively. Non-touching TFH1 cells (red borders) and B cells (blue borders) are shown schematically. Quantification of the nearest neighbor distances between TFH1 and proliferating B cells was done using Phenoptr Reports in R. The FFPE slides were scanned using Vectra Polaris and analyzed with InForm software. Statistical analysis was performed using Mann Whitney test. Each dot represents the average frequencies of 20 regions of interest from 4 T21 and 3 D21l tonsils. D) Cell-to-cell communications between B and CD4 T cells in D21 and T21 tonsils as analyzed by the CellChat program. Circle plots showing the interactions and strengths for different inflammatory pathways (IFN-II, TNF, IL6 and chemokines pathways) among different B and CD4 T clusters previously described in Figures 3 and 5.

Consistent with this relationship, spatial analysis revealed a trend toward increased proximity between TFH1-like cells (CD3⁺PD-1⁺BCL6⁺TBET⁺) and Ki67⁺ proliferating B cells in extrafollicular areas of T21 tonsils. Indeed, whereas in D21 tonsils, TFH1 like– Ki67+B cell conjugates were predominantly localized within well-organized GCs consistent with efficient selection niches, in T21 tonsils, TFH1-Ki67+ B cell aggregates are increased at the extrafollicular areas (**Fig. 6C, lower panel**). These findings support a shift in the anatomical localization of T–B interactions rather than a strict increase in their overall frequency (**Fig. 6C**).

To determine whether the altered spatial organization of T–B interactions was accompanied by changes in the underlying communication networks, we next inferred ligand–receptor interactions across immune cell populations using CellChat. Cell–cell communication analysis revealed extensive rewiring of signalling networks in T21 tonsils (**Fig. 6D**). Canonical GC–associated pathways, including ICOS, CD40, BAFF and CXCL signalling were less prominent within the inferred communication network. (**Extended Data Fig. 6B**), whereas inflammatory cytokine networks such as IFN-II, TNF, IL6 and CCL were more broadly distributed across CD4 effector/helper and memory compartments in T21 (**Fig. 6D**).

To assess the relationship between CXCR3 expression and tissue-level immune features, we performed an integrated analysis of immunophenotypic and histological parameters across individual samples. Unsupervised clustering revealed a clear segregation of T21 tonsils, characterized by broadly increased CXCR3 expression across multiple B- and T-cell subsets, including memory CD4⁺ and CD8⁺ T cells and class-switched B cells. This CXCR3-high state was associated with increased expression of activation markers, as well as variation in GC features. In contrast, D21 samples displayed lower and more heterogeneous profiles. These data indicate that T21 is associated with a coordinated, tissue-wide CXCR3-associated immune program (**Extended data Fig. 6B**).

Together, these findings indicate that T21 remodels the cellular and communication architecture of secondary lymphoid organs, shifting immune interactions away from canonical germinal center networks toward inflammatory circuits. Such tissue-level rewiring provides a mechanistic framework linking chronic interferon-associated inflammation with altered T- and B-cell differentiation and defective humoral immunity.

## Discussion

Despite decades of investigation, the mechanisms underlying immune dysregulation in T21 remain incompletely understood. Chronic interferon signaling, driven in part by triplication of four interferon receptor genes on chromosome 21, has been linked to widespread immune abnormalities in peripheral blood, including impaired B-cell maturation, skewed T-cell differentiation and dysregulated innate responses ^4,5,9,10^. Our findings extend this paradigm to SLO, suggesting that systemic immune abnormalities may arise, at least in part, from altered cellular interactions and differentiation programs within lymphoid tissue microenvironments

We identify profound remodeling of CD4⁺ T-cell differentiation, characterized by contraction of canonical TFH populations and expansion of inflammatory helper states. T21 tonsils displayed increased frequencies of TFH1-like cells producing IFNγ and IL-21 together with enrichment of interferon-responsive transcriptional programs. Because these alterations occur in the absence of acute infection, our data suggest that sustained inflammatory cues are sufficient to promote TFH1-like differentiation within human lymphoid tissues. In addition, we identified a ZEB2⁺GZMH⁺ CD4⁺ T-cell population enriched in T21, consistent with cytotoxic helper-like or age-associated helper T cells previously described in aging and autoimmune diseases ^35–37^. Together, these findings support a model in which chronic inflammatory signaling promotes inflammatory and effector differentiation at the expense of conventional follicular helper programs.

A particularly striking observation was the widespread increase in CXCR3 expression across both T-cell and B-cell compartments. Beyond serving as a marker of interferon exposure, CXCR3 regulates migration toward CXCL9, CXCL10 and CXCL11 and has been associated with inflammatory tissue positioning at the expense of CXCR5-dependent follicular localization. The concomitant increase in CXCR3 expression in TFH1-like, memory and B-cell populations therefore raises the possibility that chronic interferon signaling not only reprograms immune cell transcriptional states but also reshapes lymphoid positioning cues, promoting redistribution of adaptive immune cells away from canonical GC niches. Such a mechanism may contribute to the reduced follicular organization observed in T21 tonsils and provides a potential link between chronic inflammatory signaling and impaired follicular immunity^40–43^.

Consistent with this model, T21 tonsils exhibited reduced TFH-associated neighborhoods, smaller lymphoid follicles and profound changes in B-cell differentiation. Although the overall composition of the B-cell compartment remained largely preserved, probably due to the early age of the cohort, T21 B cells displayed enhanced interferon- and activation-associated transcriptional programs together with reduced proliferative signatures. Notably, we observed accumulation of PRE_GC cells accompanied by altered clonal connectivity and reduced GC-associated neighborhoods. Rather than simply reflecting increased activation, these findings suggest that B-cell differentiation is diverted before completion of canonical GC maturation, leading to accumulation of transitional pre-GC states and increased reliance on alternative differentiation pathways. Such a bottleneck in GC progression is consistent with the reduced proliferative programs, diminished follicular architecture and impaired humoral responses previously described in DS ^14–18^.

Our clonal analyses further support this interpretation. T21 tonsils displayed broader clonal connectivity linking ACT, cycling, pre-GC, GC and memory populations, together with expansion of PRE_GC clonotype sets. Rather than representing isolated differences in cell abundance, these findings suggest coordinated alterations in lineage commitment and cell fate decisions. Similarly, T-cell trajectory analyses demonstrated diversion away from canonical TFH differentiation toward inflammatory helper and cytotoxic states. Together, these observations support a model in which chronic interferon signaling redirects both T- and B-cell differentiation away from highly organized follicular programs toward inflammatory trajectories.

The CD8⁺ compartment was similarly remodeled. T21 tonsils displayed increased inflammatory and cytotoxic programs together with enrichment of resident memory-like states, in the absence of transcriptional features associated with classical exhaustion. These observations argue against chronic antigen overstimulation as the primary driver and instead support sustained inflammatory instruction within lymphoid tissues. Mechanistically, altered interactions between CD8⁺ T cells and DCs may further contribute to tissue reorganization. We observed depletion of XCL1⁺XCL2⁺ cytokine-enriched CD8⁺ T cells, previously implicated in recruitment and positioning of XCR1⁺ dendritic cells and the organization of cross-presenting niches^44^. This defect may be compounded by expansion of CD56⁺ dendritic cells, a population associated with inflammatory activation and enhanced interactions with cytotoxic lymphocytes, potentially contributing to disruption of canonical cDC1 niches ^33,34^.

Collectively, these findings support a model in which chronic interferon-biased inflammation caused by T21 remodels the immune microenvironment of SLOs at multiple levels. Beyond inducing inflammatory transcriptional programs, sustained interferon signaling appears to alter chemokine responsiveness, tissue positioning and lineage commitment across adaptive immune populations. By simultaneously promoting CXCR3-associated inflammatory migration and diverting differentiation away from canonical follicular programs, chronic inflammatory signaling observed in T21 tonsils, may impair GC organization and compromise the generation of high-quality humoral immunity

More broadly, our findings suggest that chronic interferon signaling may represent a general mechanism by which persistent inflammatory cues reshape lymphoid tissue organization and adaptive immune cell fate decisions. Similar immune remodeling has been described in autoimmune diseases such as systemic lupus erythematosus, where dysregulated GC responses, expansion of CXCR3⁺ inflammatory lymphocytes and increased extrafollicular differentiation accompany defective humoral immunity^22,23,45^. Our data extend these observations by demonstrating that chronic interferon exposure can simultaneously reshape tissue architecture, migratory programs and differentiation trajectories within T21 SLO and also probably, in other similar pathophysiological human situations.

Several limitations should be considered. The relatively small number of samples included in the single-cell analyses may limit assessment of inter-individual variability, and tonsils represent specialized SLO continuously exposed to environmental antigens. Furthermore, direct assessment of affinity maturation, antibody quality and memory responses will be necessary to establish the functional consequences of the altered GC dynamics described here. Nevertheless, our findings provide a tissue-level framework that may help explain recently described defects in circulating memory B-cell compartments and humoral immunity in individuals with T21 ^14–18^.

In summary, our study identifies chronic interferon-biased inflammation as a major determinant of immune cell differentiation and spatial organization within SLO in T21. We propose a model in which sustained interferon signaling reshapes both immune cell positioning and lineage commitment by altering chemokine responsiveness, follicular residency and differentiation trajectories. This coordinated immune rewiring redirects adaptive immune responses away from canonical GC programs toward inflammatory states, providing a mechanistic framework linking chronic interferon signaling with defective humoral immunity in DS and potentially in other interferon-driven inflammatory disorders.

## Methods

### Sample Donors and Ethics

Palatine tonsils were obtained at Clínica COAT, SRL (Centro Oto Audiologico de Alta Tecnologia Clinica), Córdoba, Argentina. from pediatric donors undergoing tonsillectomy for obstructive hypertrophy or obstructive sleep apnea. Written informed consent was obtained from parents or legal guardians prior to sample collection. The study protocol was approved by the local institutional ethics committee (REPIS #3831) and conducted in accordance with the Declaration of Helsinki and international ethical guidelines.

The study compared children with T21 and age- and sex-matched euploid controls (D21). Strict exclusion criteria were applied to minimize confounding variables. Tonsils from 17 T21 donors and 8 D21 controls were processed.

Clinical and demographic characteristics are summarized in **Table 1**. The complete cohort was used for flow cytometry analyses. A subset of samples (D21, n=3; T21, n=3) was selected for single-cell multi-omics analyses, and a partially overlapping subset was used for multiplex immunofluorescence analyses **(Supplementary Table 1)**

**Table 1.** Clinical characteristics of study participants.

| **Variable** | **D21 (Controls)**<br>**(n = 8)** | **T21 (Down syndrome)**<br>**(n = 17)** |
| --- | --- | --- |
| **Age, years (mean ± SD)** | 6.0 ± 2.6 | 6.9 ± 3.2 |
| **Sex, n (%)** |  |  |
| **Male** | 4 (50.0%) | 10 (58.8%) |
| **Female** | 4 (50.0%) | 7 (41.2%) |
| **Trisomy diagnosis, n (%)** | N/A |  |
| **Complete trisomy 21** | N/A | 17/17 (100.0%) |
| **Partial trisomy** | N/A | 0/17 (0.0%) |
| **Mosaicism** | N/A | 0/17 (0.0%) |
| **Unspecified** | N/A | 0/17 (0.0%) |
| **Institutionalized, n (%)** | 0/8 (0.0%) | 0/17 (0.0%) |
| **Clinical history, n (%)** |  |  |
| **Leukemia** | 0/8 (0.0%) | 1/17 (6.7%) |
| **Congenital heart defects (CHD)** | 0/8 (0.0%) | 1/17 (6.7%) |
| **Autoimmune conditions**<br>**(celiac disease,**<br>**hypothyroidism,**<br>**alopecia areata, atopic**<br>**dermatitis/eczema,**<br>**vitiligo, and psoriasis)** | 0/8 (0.0%) | 3/17 (20.0%) |

*Abbreviations: CHD, congenital heart disease; D21, euploid controls; T21, trisomy 21.*

### Tissue processing

Tonsillar mononuclear cells (TMCs) were obtained from non-infected tonsils collected during clinically indicated tonsillectomies at COAT. All samples were processed within 3 hours after surgery. Tonsils were transported on ice in RPMI-1640 medium (Gibco, cat. 31800089) supplemented with 10% fetal bovine serum (FBS, NATOCOR), 1% GlutaMAX (Gibco, cat. 25030081), and 1% Penicillin-Streptomycin (Gibco, cat. 15140122). Tissues were weighed and washed with PBS and mechanically dissociated using sterile scalpels or scissors. When two tonsils from the same donor were available, one was allocated for histological analysis and the other for cell isolation.

Tissue fragments were centrifuged, resuspended in digestion medium containing 0.1 mg ml⁻¹ of Liberase TL (Roche) and 0.1 mg ml⁻¹ of DNase I (Sigma Aldrich), and incubated for 40–45 min at room temperature with agitation. Enzymatic digestion was stopped with EDTA (Gibco) at a final concentration 5 mM. Cell suspensions were filtered through 100μm sterile cell strainers (Corning) and mononuclear cells were isolated by density-gradient centrifugation using Ficoll-Paque PLUS (Sigma Aldrich). Cell purity consistently exceeded 95%.

Cells were counted by trypan blue exclusion and cryopreserved at 20–50 × 10⁶ cells per vial in RPMI containing 10% FBS and 10% DMSO using controlled-rate freezing, followed by storage in liquid nitrogen.

### Flow cytometry

#### Sample preparation and staining

Cryopreserved TMCs were rapidly thawed, washed, and plated at 1 × 10⁶ cells per well in 96-well U-bottom plates (Greiner Bio-One). Surface staining was performed using fluorochrome-conjugated monoclonal antibodies specific for lineage, activation, and differentiation markers **(Supplementary Tables 2-5).**

For chemokine receptor detection, cells were incubated with anti-CXCR3 and anti-CXCR5 antibodies at 37 °C for 20 min prior to additional surface staining at 2–8 °C. Viability was assessed using LIVE/DEAD Fixable Aqua dye. When transcription factor or intracellular protein detection was required, cells were fixed and permeabilized using the Foxp3/Transcription Factor Staining Buffer Set (Invitrogen) following the manufacturer’s instructions, and stained with the appropriate antibody cocktails **(Supplementary Tables 2-5).**

#### Intracellular cytokine staining

For functional assays, TMCs were stimulated with phorbol 12-myristate 13-acetate (50 ng ml⁻¹), ionomycin (1 μg ml⁻¹), and a protein transport inhibitor cocktail for 4 h at 37 °C. Cells were then stained for viability and surface markers, fixed, permeabilized, and stained for intracellular cytokines using the antibodies detailed in **Supplementary Table 3**.

### Data acquisition

Data were acquired using a BD LSRFortessa™ (BD Biosciences) multicolor flow cytometer and analyzed using FlowJo™ v10.6.2 software (BD Bioscience) and R-based computational workflows.

### Omics analysis

#### Cell sorting

For transcriptomic analyses, TMCs from 3 T21 and 3 D21 donors were stained with a viability dye and an anti-CD19 antibody and sorted using a high-speed cell sorter. CD19⁻ and CD19⁺ fractions were recombined at a 70:30 ratio to preserve T-cell representation while maintaining B-cell diversity. The relative abundance of immune populations was quantified by flow cytometry prior to sorting (See **Fig. 1D**).

#### CITE-seq and TCR-seq library preparation

Mixed sorted fractions from the 6 donors were loaded onto the Chromium Controller (10x Genomics). Single-cell gene expression, paired TCR sequencing, and surface protein profiling were performed using the Chromium GEM-X Single Cell 5′ v3 Gene Expression with Feature Barcoding technology for Cell Surface Protein (10x Genomics), according to the manufacturer’s instructions. Cell surface proteins were labeled using the TotalSeq™-C Human Universal Cocktail, V1.0 (BioLegend, Cat. No. 399905). Libraries were sequenced on an Illumina NovaSeq 6000 platform.

#### CITE-seq data analysis

Base call (BCL) files were demultiplexed and converted to FASTQ format using bcl2fastq2 (v2.20). Gene expression and antibody-derived tag (ADT) count matrices were generated using Cell Ranger (10x Genomics) with the GRCh38 (hg38) reference genome. Downstream analyses were performed in R using Seurat (v4.3.0.1).

Cells with fewer than 200 or more than 5,000 detected genes, fewer than 700 or more than 30,000 unique molecular identifiers (UMIs), or more than 25% mitochondrial transcripts were excluded to remove low-quality cells, potential doublets, and dying cells. Y chromosome genes were excluded from downstream analyses to minimize clonal overrepresentation, sex-related bias, and technical noise.

RNA expression data were normalized using the LogNormalize method implemented in NormalizeData. Highly variable genes were identified using FindVariableFeatures with the vst selection method, followed by data scaling with ScaleData and principal component analysis (PCA).

For the CD4⁺ T cell subset analysis, TCR- and BCR-related genes (identified by the gene symbol patterns ^TR[ABGD][CVJ] and ^IG[HKL][CVDJM], respectively) were additionally excluded from the RNA assay prior to normalization, together with Y chromosome-linked genes, to further reduce clonal and sex-related bias. Batch effects across patients were then corrected on the RNA PCA embedding using Harmony (RunHarmony, group.by.vars = “patient”), and the resulting Harmony-corrected embedding was used in place of the standard PCA for downstream multimodal integration.

For ADT data, all measured protein features were retained for downstream analyses. ADT counts were normalized using centered log-ratio (CLR) normalization (margin = 2), scaled, and subjected to PCA.

RNA and ADT modalities were integrated using the weighted nearest neighbor (WNN) approach implemented in FindMultiModalNeighbors, using the first 50 RNA principal components and the first 30 ADT principal components. Cell clustering was performed on the weighted shared nearest neighbor (wsnn) graph using the Smart Local Moving (SLM) algorithm (algorithm = 3) with a resolution of 4.

Uniform Manifold Approximation and Projection (UMAP) was generated based on the weighted nearest neighbor graph for visualization. To determine DEGs between clusters, the FindAllMarkers function and the Wilcoxon rank sum test were used and an adjusted p-value < 0.05. Neighbourhood-based differential abundance (DA) testing was performed using Milo (v1.7.0) by embedding cells in a Harmony-corrected k-NN graph and modeling neighbourhood counts via a Negative Binomial GLM ^46^.

Pathway enrichment analyses were conducted using clusterProfiler with MSigDB Hallmark gene sets^47^ and Blood Transcription Modules (BTM)^32^. Cell–cell communication inference was carried out using CellChat (v1.6.1) ^48^.

#### Velocity and trajectory analysis

Cell embeddings and cluster annotations from the integrated Seurat object (based on WNN analysis of RNA and protein [ADT] modalities) were used to compute a diffusion map with the destiny R package (v3.20.0), based on the first 50 Harmony-corrected principal components, retaining 5 diffusion components (n_eigs = 5, density_norm = TRUE). The first two diffusion components (DC1, DC2) were used as a two-dimensional embedding. Raw counts, Harmony embeddings, WNN-UMAP coordinates, and diffusion map coordinates were exported from the Seurat object and reconstructed as an AnnData object in Python using scanpy and anndata. TCR-related genes (TRAV, TRAC, TRAJ, TRBV, TRBC, TRBJ, TRBD) were excluded prior to downstream analysis. RNA velocity was analyzed separately for each condition (D21 and T21). For each condition, spliced and unspliced count matrices were generated per patient from the corresponding CellRanger BAM files using velocyto (v0.17.17), producing one loom file per patient; individual loom files were concatenated and merged with the AnnData object using scVelo’s merge function. Data were filtered and normalized with scVelo’s (v0.3.4)^49^. filter_and_normalize function, and first- and second-order moments were computed using the top 50 principal components (Harmony-corrected) and 30 nearest neighbors. RNA velocity was estimated independently for each condition using the stochastic model of transcriptional dynamics and projected onto the pre-computed diffusion map embedding. Velocity confidence was calculated to assess the coherence of velocity vectors across neighboring cells^49^.

#### Single-cell TCR-seq data analysis

FASTQ files were processed using the cellranger vdj pipeline with GRCh38/hg38 as the reference genome. The resulting filtered_contig_annotations.csv files were imported into R and processed per patient. Contigs were filtered to retain high-confidence, productive TRB (beta chain) sequences (high_confidence = TRUE, productive = TRUE, chain = “TRB”); for cells with more than one *TRB* contig, the dominant contig was selected based on the highest UMI count (ties broken by read count). Clonotype identity was defined as the combination of *TRB V* gene, *J* gene, CDR3 amino acid sequence, and CDR3 nucleotide sequence. Only cells with a valid, unambiguous *TRB* clonotype assignment were retained for downstream clonal analyses.

#### Single-cell BCR-seq data analysis

BCR sequencing data were processed analogously to the TCR data described above. High-confidence, productive immunoglobulin heavy chain (*IGH*) contigs were retained per patient, and the dominant *IGH* contig per cell was selected by UMI count (ties broken by read count). B-cell clonotype identity was defined as the combination of *IGH V, D*, and *J* gene usage together with the CDR3 nucleotide and amino acid sequences. Only cells with an unambiguous *IGH* clonotype assignment were retained for downstream clonal analyses.

#### Clonal overlap analysis

Clonal overlap between cell clusters was quantified using the Morisita-Horn Index (MHI)^50^, calculated for each pairwise combination of clusters using clonotypes with more than one cell (expanded clones) that were shared across at least two clusters. Statistical significance was assessed by a permutation test: for each pair of clusters, clone-abundance vectors were independently reshuffled 1,000 times to generate a null distribution of the index, and an empirical p-value was calculated as the proportion of permuted values greater than or equal to the observed index. This analysis was performed on the T21 and D21 groups.

Clonotype sharing across the same set of clusters was additionally visualized using UpSet plots (ComplexHeatmap, v2.22.0), generated separately for the D21 and T21 groups, showing the size of each cluster-intersection set of shared *TRB* clonotypes.

#### Clonal expansion in trajectory space

Expanded TRB clonotypes (≥2 cells) identified in CD4⁺ T cells were projected onto the diffusion map embedding described above (RNA velocity and trajectory analysis section). For each expanded clonotype, a centroid was calculated as the mean DC1/DC2 coordinate across all cells belonging to that clone, colored by main cluster (TFH, CM_preTFH, CM_CXCR3, Trans_mem, T_helper, or CTL) and sized proportionally to clone size; non-expanded and unassigned cells were displayed in the background. Plots were generated separately for the D21 and T21 groups.

#### Gene signature scoring

Cell-intrinsic transcriptional signatures were quantified using Seurat’s AddModuleScore function with default parameters. For CD4⁺ T cells, gene sets corresponding to the Hallmark Interferon Gamma Response and Hallmark Allograft Rejection signatures were obtained from the Molecular Signatures Database (MSigDB, Hallmark collection) using the msigdbr package (v10.0.2)^47^ and filtered to retain genes detected in the dataset. For B cells, activation and proliferation signatures were defined using manually curated marker gene sets (activation: immediate-early and NF-κB-responsive genes including FOS, JUN, EGR1–3, CD69, CD83, CD86, NFKBIA, and NR4A1–3; proliferation: cell-cycle and mitotic genes including MKI67, TOP2A, PCNA, MCM2–7, and CCNB1/2), likewise filtered to genes present in the dataset. Module scores were compared between D21 and T21 conditions within each cell subtype/cluster (cluster_TFH for CD4⁺ T cells; subtypes for B cells) using violin plots, with the group median indicated and statistical significance assessed by two-sided Wilcoxon rank-sum test.

### Multiplex immunohistochemistry and immunofluorescence

#### Tissue processing

Tonsil specimens were fixed in 10% (v/v) neutral buffered formalin (pH 7; Biopur), dehydrated through a graded ethanol series, cleared in xylene (Cicarelli), and embedded in paraffin (Cicarelli) at 60°C. Formalin-fixed, paraffin-embedded (FFPE) tissue blocks were sectioned at 5 μm using a rotary microtome (Thermo Scientific HM 325). Sections were floated on a 45°C water bath to remove wrinkles and mounted onto glass slides for subsequent processing.

#### Follicle size quantification and correlation analysis

Lymphoid follicle size was quantified using multispectral immunofluorescence images acquired with the PhenoImager Fusion system (Akoya Biosciences) and analyzed in QuPath to quantify BCL-6 (Abcam, cat. ab249707) positive follicular areas (10 to 35 follicles per slide). Follicular area and follicle density (number of follicles per cm²) were compared between experimental groups and visualized as violin plots in R (D21, n=6; T21, n=15).

To investigate the relationship between lymphoid architecture and T cell composition, follicle size measurements obtained with QuPath were integrated with flow cytometry data from matched samples. The mean follicle diameter for each sample was estimated as the square root of the measured follicular area and correlated with the frequencies of T follicular helper (TFH), defined as CXCR5hi PD1hi CXCR3neg (expressed as a percentage of live cells), and CM_CXCR3, defined as CD4⁺PD1⁻CXCR3⁺ T cells (excluding naïve and regulatory T cells). Pearson correlation coefficients were calculated in R, and correlations were visualized as scatter plots with fitted linear regression lines and 95% confidence intervals. Correlation coefficients (R) and corresponding P values were reported for each analysis (D21, n=5; T21, n=12).

#### Multiplex immunofluorescence (mIF)

Sequential Opal-based multiplex immunofluorescence was performed on paraffin sections to detect nine markers simultaneously (D21, n=3; T21, n=4). Antibody panels are listed in **Supplementary Table 6**. Each staining cycle included antigen retrieval, blocking, primary antibody incubation, HRP-conjugated secondary antibody incubation, fluorophore deposition, and antibody stripping prior to the next cycle. Slides were scanned using multispectral imaging systems, and regions of interest corresponding to T-cell and B-cell zones were selected.

#### Image analysis and spatial statistics

Spectral unmixing, tissue segmentation, and single-cell phenotyping were performed using supervised machine-learning algorithms. Quantitative analyses included cell density, relative frequencies, spatial proximity within a 15 μm radius, and minimum intercellular distances between defined phenotypes. Quantification of the nearest neighbor distances between cells was done using Phenoptr Reports in R. The FFPE slides were scanned using Vectra Polaris and analyzed with InForm software. Statistical analysis was performed using Mann Whitney test.

#### Use of AI Tools

Large language models, including ChatGPT (OpenAI, GPT-4) and Claude (Anthropic), were used to assist in refining the wording, grammar, and clarity of sections of the manuscript, including the introduction and methods. All scientific content, analysis, and conclusions were conceived, conducted, and validated by the authors.

## Data availability

Data availability: Raw and processed single-cell RNA-seq, single-cell TCR-seq, and single-cell CITE-seq data are available under accession number GSE3XXXX

## Acknowledgements

We are grateful to Drs Santiago Boccardo, Paula Abadie, Pilar Crespo, Maria Soledad Miró, Luciana Reyna, Paula Icely for their technical support in sample handling, flow citometry and histotechnology at CIBICI-CONICET. We also thank the Functional Genomics Center Zurich (University of Zurich) and the Cytometry Facility (University of Zurich) for their technical support.

## Author Contributions

JD, MM and NN designed the study and analysed the data; JD and JB performed the experiments; LB, PA, SD, AFG, PB and REB contributed in the sample processing, flow cytometry, data analysis and in other experiments; JCK and VC performed the sorting experiments; CV and CCM selected the patients, JTB and WR contributed in the data analysis; BB, JME, JTB and EP carefully read the manuscript, JD, MM and NN wrote the manuscript.

## Legends for the Supplementary Figures

**Extended Data Figure 1.**
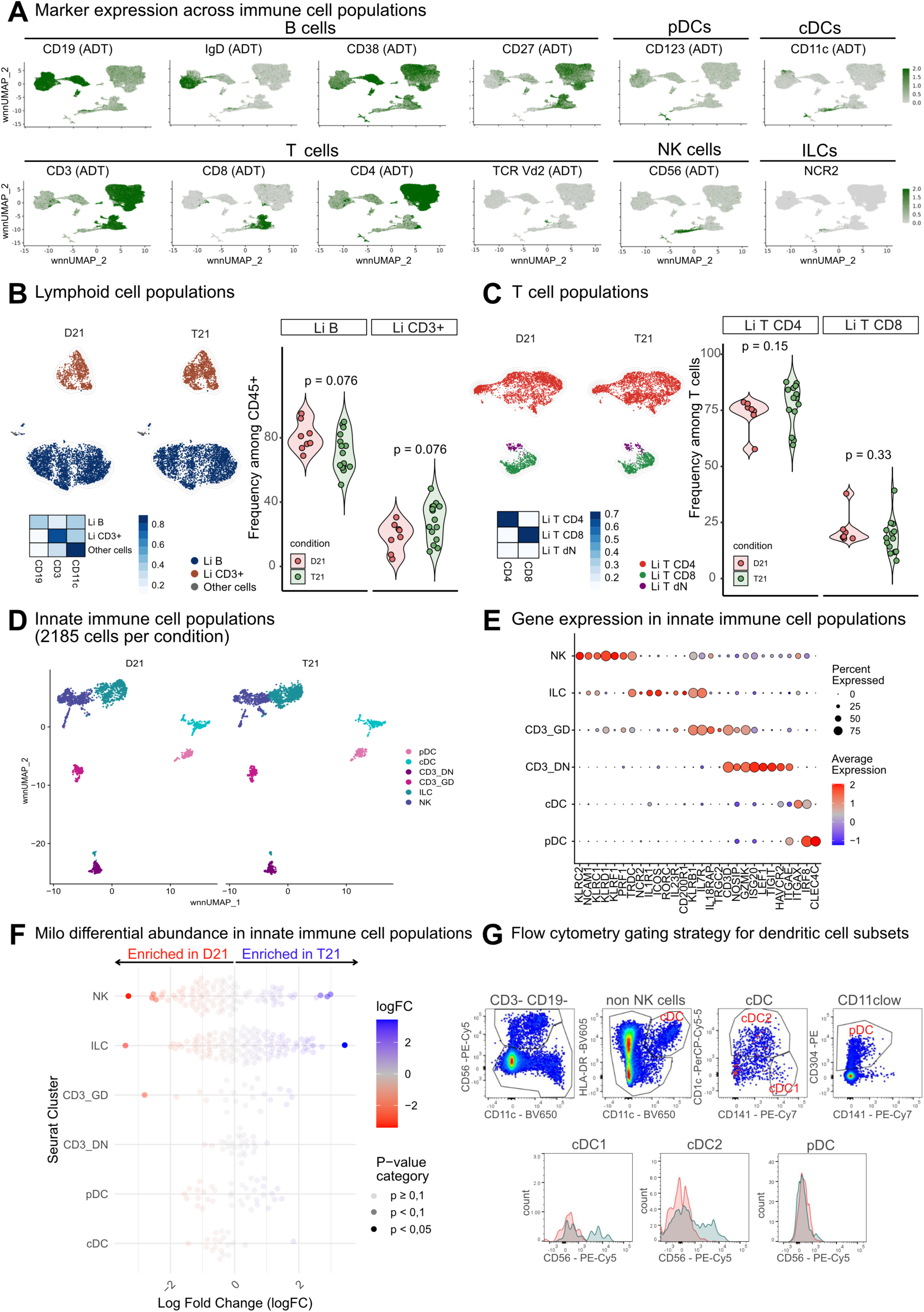
Annotation and validation of tonsillar immune populations identified by multimodal single-cell profiling. **(A)** Representative antibody-derived tag (ADT) and transcriptomic markers used for annotation of major immune populations in the CITE-seq dataset. Major lymphoid and dendritic cell populations were primarily identified using surface protein expression, including markers for B cells (CD19, IgD, CD27 and CD38), T cells (CD3, CD4 and CD8), pDCs (CD123), cDCs (CD11c), NK cells (CD56) and γδ T cells (TCRVδ2). ILCs were identified based on the expression of canonical transcriptional markers owing to the absence of unique surface markers in the ADT panel (i.e NCR2 as shown in the figure) . **(B)** FlowSOM/UMAP representation of flow cytometry data showing the distribution of major lymphoid populations, including B cells, CD3⁺ cells and remaining CD45⁺ populations, in D21 and T21 tonsils. The accompanying heatmap depicts the expression intensity of markers used for population annotation. Violin plots show the frequencies of B cells and CD3⁺ T cells among total CD45⁺ cells in the validation cohort (D21, n = 8; T21, n = 15). Each dot represents one individual. P values were calculated using two-tailed Mann–Whitney tests. **(C)** FlowSOM/UMAP representation of T-cell populations identified by flow cytometry, including CD4⁺, CD8⁺ and double-negative T cells (CD3⁺CD4⁻CD8⁻), in D21 and T21 tonsils. The accompanying heatmap shows the expression of markers used for subset annotation. Violin plots depict the frequencies of each subset among total CD3⁺ T cells in the validation cohort. Each dot represents one individual. P values were calculated using two-tailed Mann–Whitney tests. **(D)** wnnUMAP representation of innate immune populations extracted from the global dataset, including pDC, cDC, NK cells, ILC, and CD3_DN T cells. The accompanying dot plot displays representative transcriptomic markers used for cluster annotation **(E)**. **(F)** Milo differential abundance analysis of innate immune populations comparing T21 and D21 tonsils. Beeswarm plot showing the distribution of neighborhood log fold changes (logFC) across annotated innate immune clusters. Each point represents a transcriptional neighborhood assigned to a given cell population and is positioned according to its differential abundance between conditions. Point color indicates the magnitude and direction of abundance changes (red, enriched in D21; blue, enriched in T21), while point size denotes statistical significance categories based on adjusted P values. Point size reflects the statistical significance category based on the P-value **(G)** Representative flow cytometry gating strategy used for the identification of dendritic cell subsets. Sequential gating of CD3⁻CD19⁻ non-NK cells allowed discrimination of cDC1, cDC2 and pDC populations based on the expression of HLA-DR, CD11c, CD141, CD1c and CD304. Representative histograms illustrate CD56 expression across dendritic cell subsets.

**Extended Data Figure 2.**
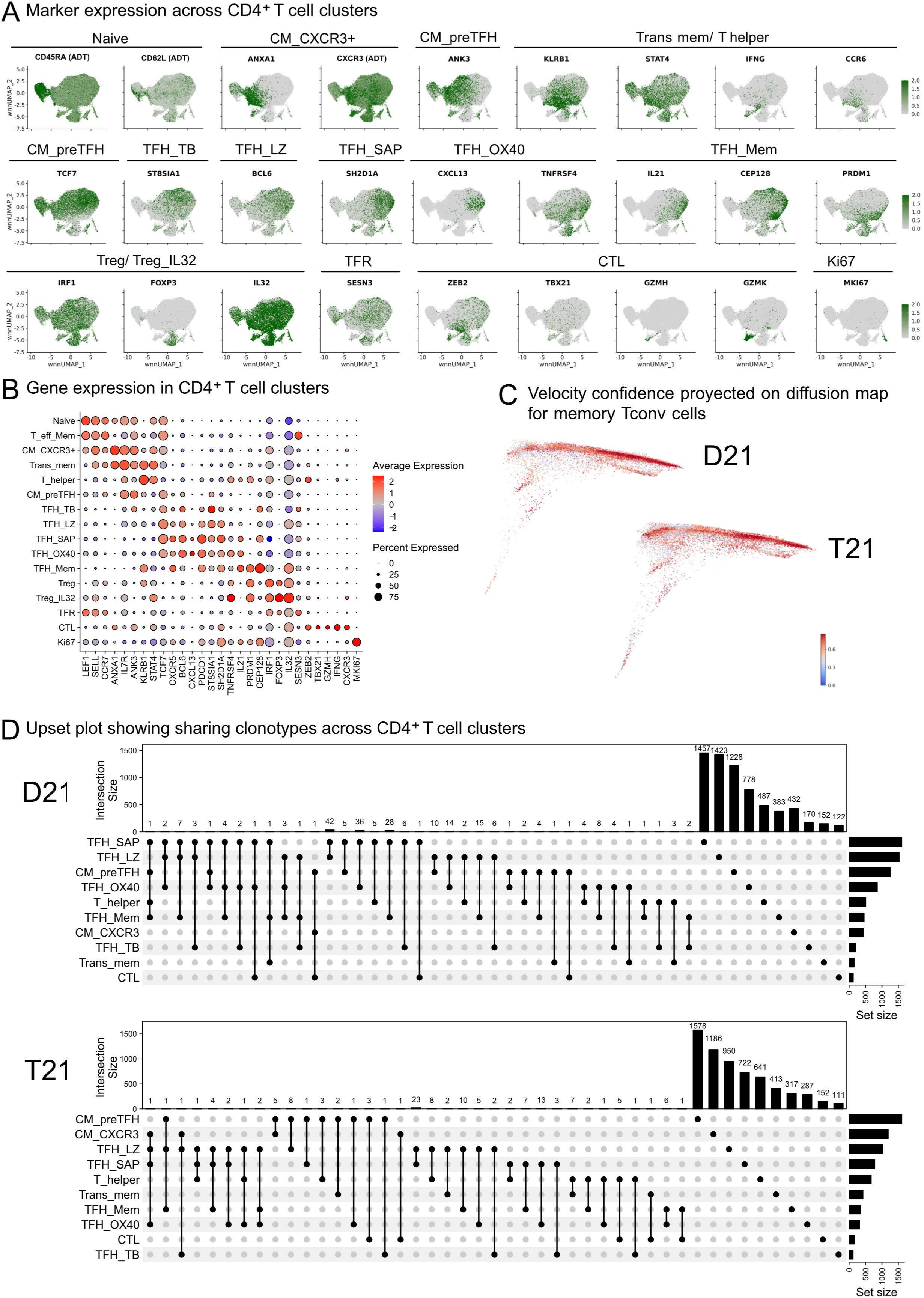
Characterization of tonsillar CD4⁺ T-cell states and clonotype relationships in T21 and D21 tonsils. (A) Representative marker expression across CD4⁺ T-cell clusters identified by CITE-seq. UMAP projections show the expression of representative antibody-derived tags (ADT) and transcriptomic markers used for cluster annotation, including naïve markers (CD45RA, CD62L), memory-associated markers (ANXA1, CXCR3, ANK3, KLRB1, STAT4), canonical TFH markers (TCF7, BCL6, CXCL13, PDCD1, IL21, SH2D1A and TNFRSF4), regulatory markers (FOXP3, IL32, SESN3), cytotoxic markers (ZEB2, TBX21, GZMH and IFNG), and proliferation markers (MKI67). (B) Dot plot summarizing the expression of representative genes across annotated CD4⁺ T-cell populations and clusters. Dot size indicates the proportion of expressing cells, whereas color intensity represents scaled average expression levels. The identified populations include naïve, memory, TFH, regulatory, cytotoxic and proliferating CD4⁺ T-cell states. (C) RNA velocity confidence scores projected onto diffusion map embeddings of memory CD4⁺ T cells from D21 and T21 tonsils. Color intensity represents the confidence of velocity estimates and supports the reliability of inferred transcriptional trajectories across cellular states. (D) UpSet plots showing TCR clonotype sharing across CD4⁺ T-cell populations in D21 and T21 tonsils. Vertical bars indicate the number of clonotypes shared among the indicated populations, whereas horizontal bars represent the total clonotype content of each subset. Numbers on top of the vertical bars indicate the number of shared clonotypes. Set sizes for each population are shown on the right.

**Extended Data Figure 3.**
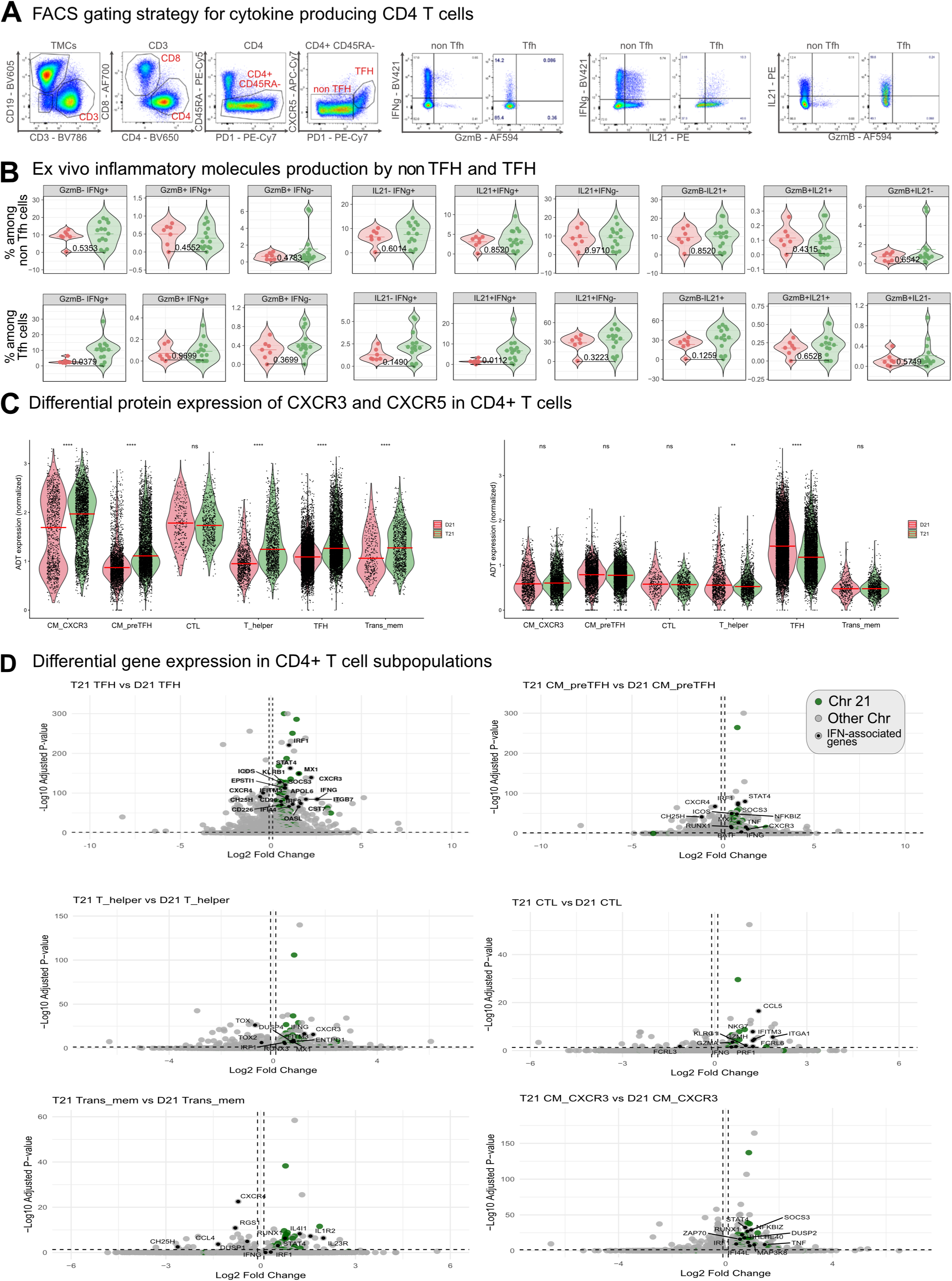
Functional and transcriptional characterization of interferon-biased CD4⁺ T-cell remodeling in T21 tonsils. **(A)** Flow cytometry gating strategy used to identify cytokine-producing CD4⁺ T-cell populations. Representative gating strategy for the identification of total CD4⁺ T cells, TFH (PD1^hiCXCR5^hi) and non-TFH populations, followed by assessment of intracellular IFNγ, IL-21 and granzyme B expression after ex vivo stimulation. **(B)** Ex vivo inflammatory molecule production by TFH and non-TFH populations. Frequencies of IFNγ-, IL21- and granzyme B-producing cells among non-TFH (upper panels) and TFH populations (lower panels) from D21 and T21 tonsils. Each dot represents one individual sample. P values were calculated using two-tailed Mann– Whitney tests. **(C)** Differential protein expression of CXCR3 and CXCR5 across CD4⁺ T-cell states. Quantification of normalized ADT expression of CXCR3 (CD183) and CXCR5 (CD185) across major CD4⁺ T-cell populations. **(D)** Differential gene expression across major CD4⁺ T-cell populations in T21 and D21 tonsils. Volcano plots showing genes differentially expressed between T21 and D21 samples within TFH, CM_preTFH, T_helper, CTL, Trans_mem and CM populations. Green dots indicate chromosome 21-encoded genes. Multiple populations exhibited upregulation of interferon-associated genes including *IFNG, IRF1, IFITM3, MX1, STAT4, CXCR3* and *RUNX1*, whereas CTL cells preferentially upregulated cytotoxic molecules including *CCL5, NKG7, GZMH* and *PRF1*.

**Extended Data Figure 4.**
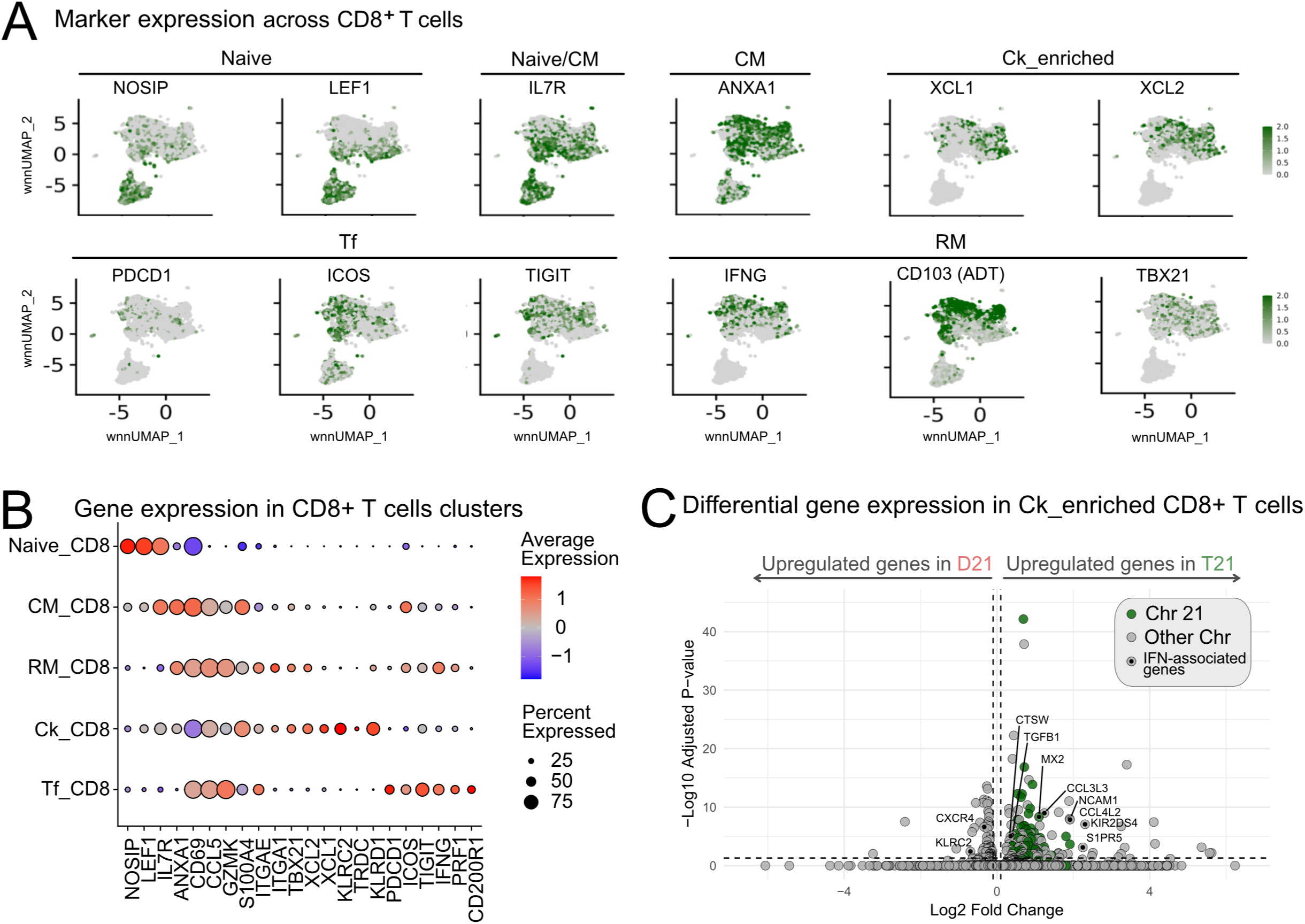
Trisomy 21 reshapes the tonsillar CD8⁺ T-cell compartment through expansion of inflammatory resident memory-like programs and depletion of cytokine-enriched states. **(A)** Marker expression across CD8⁺ T-cell populations showing representative transcriptomic and ADT markers used for cluster annotation, including naïve markers (NOSIP, LEF1, IL7R), T follicular markers (PDCD1, ICOS, TIGIT), resident-memory markers (IFNG, ITGAE/CD103, TBX21), and cytokine-enriched markers (XCL1 and XCL2). **(B)** Dot plot summarizing gene expression across annotated CD8⁺ T-cell populations. Dot size represents the fraction of expressing cells and color intensity indicates scaled average expression. The Ck_enriched population was characterized by expression of *XCL1*, *XCL2*, *KLRC2, KLRD1* and *TRDC*, whereas RM_CD8 cells preferentially expressed I*FNG, ITGAE, TBX21* and cytotoxic effector molecules. **(C)** Differential gene expression analysis of Ck-enriched CD8⁺ T cells comparing T21 and D21 tonsils. T21 Ck-enriched CD8⁺ T cells showed increased expression of chemokine- and NK-associated genes, including *CCL3L3, CCL4L2, NCAM1* and *KIR2DS4*, together with increased *TGFB1* and reduced *CXCR4* expression,

**Extended Data Figure 5.**
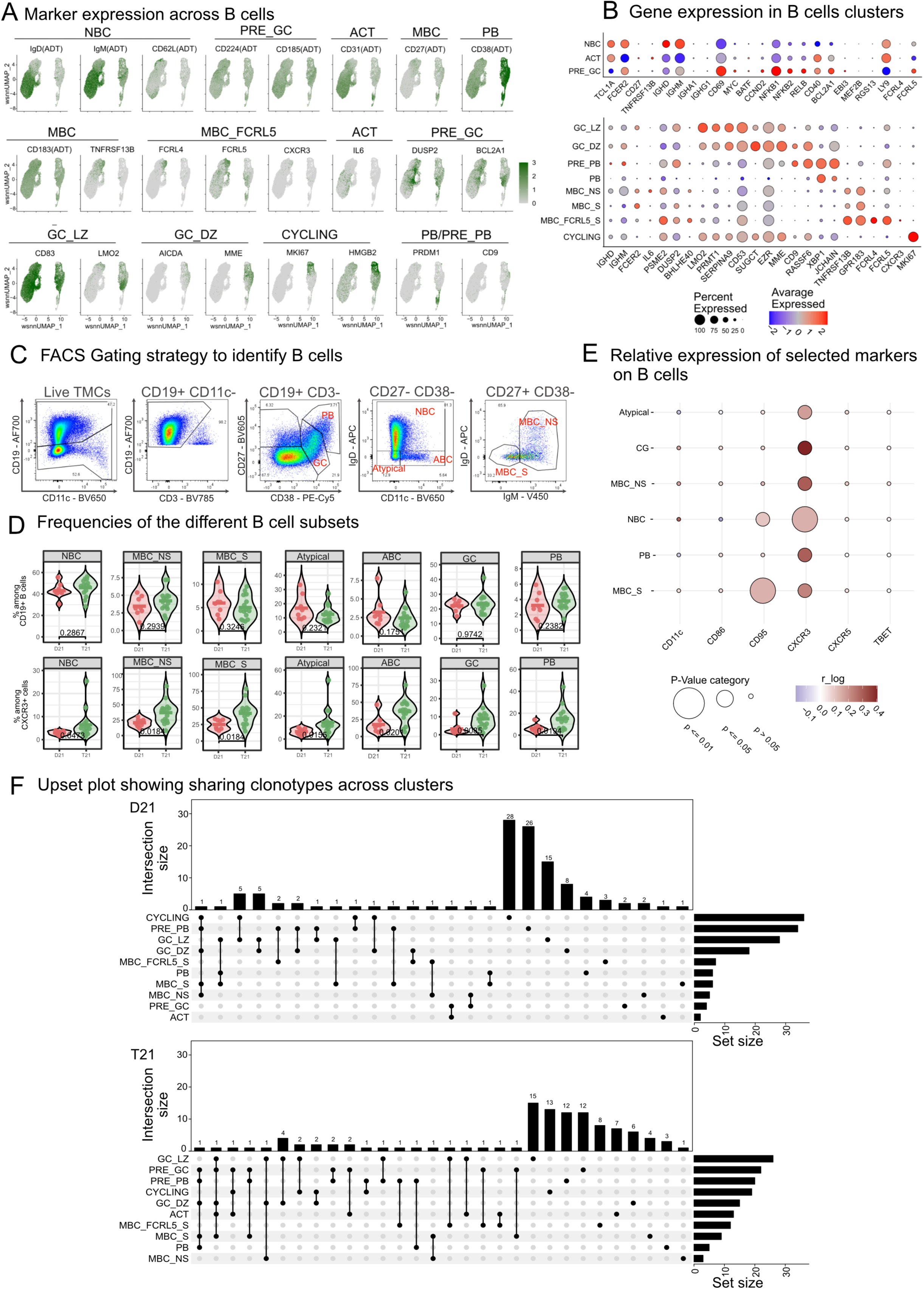
Annotation, phenotypic characterization and validation of B-cell populations in T21 and D21 tonsils. **(A)** wnnUMAP representation of B cells colored by antibody-derived tag (ADT) and gene expression levels for selected markers, including CD39, CD62L, CD49d, CD57, CXCR5 (CD185), CD49f, CD224, CD42b, CD16, CD27, *IGLL5, TNFRSF13B, FCRL4, FCRL5, TNF, NFKB1, BCL2A1, CD83, LMO2, AICDA, MME, MKI67, POLA1, HMGB2, PRDM1, and CD9.* Color intensity indicates normalized expression levels. **(B)** Dot plot summarizing gene expression across annotated B-cell populations. Dot size represents the fraction of expressing cells and color intensity indicates scaled average expression. Representative genes used for annotation include *IGHD, IGHM, FCER2, BHLHE40, SERPINA9, TNFRSF13B, FCRL4, FCRL5, MKI67, AICDA, PRDM1* and *JCHAIN*. **(C)** Representative flow cytometry gating strategy used to identify major B-cell subsets in tonsils, including naïve B cells (NBC), IgM memory B cells (MBC_NS), switched memory B cells (MBC_S), atypical B cells (Atypical), age associated B cells (ABC), germinal center B cells (GC) and plasmablasts/plasma cells (PB). **(D)** Flow cytometric quantification of major B-cell subpopulations in D21 and T21 tonsils. Upper panels show the frequency of NBC, MBC_NS, MBC_S, Atypical, ABC,GC and PB subsets among total CD19⁺ B cells. Lower panels depict the frequency of CXCR3⁺ cells within each corresponding B-cell subset. Each dot represents one individual sample. P values were calculated using two-tailed Mann–Whitney tests. **(E)** Bubble Plot showing the relative expression of selected markers across manually gated populations. Bubble size denotes statistical significance and color indicates log fold change between conditions. P values were calculated using the Mann–Whitney– Wilcoxon test . **(F)** UpSet plots showing BCR clonotype sharing across B-cell populations in D21 and T21 tonsils. Vertical bar plots indicate intersection sizes and the matrix below depicts clonotype sharing among annotated populations. Numbers on top of the bars indicate the number of shared clonotypes. Set sizes for each population are shown on the right.

**Extended Data Figure 6.**
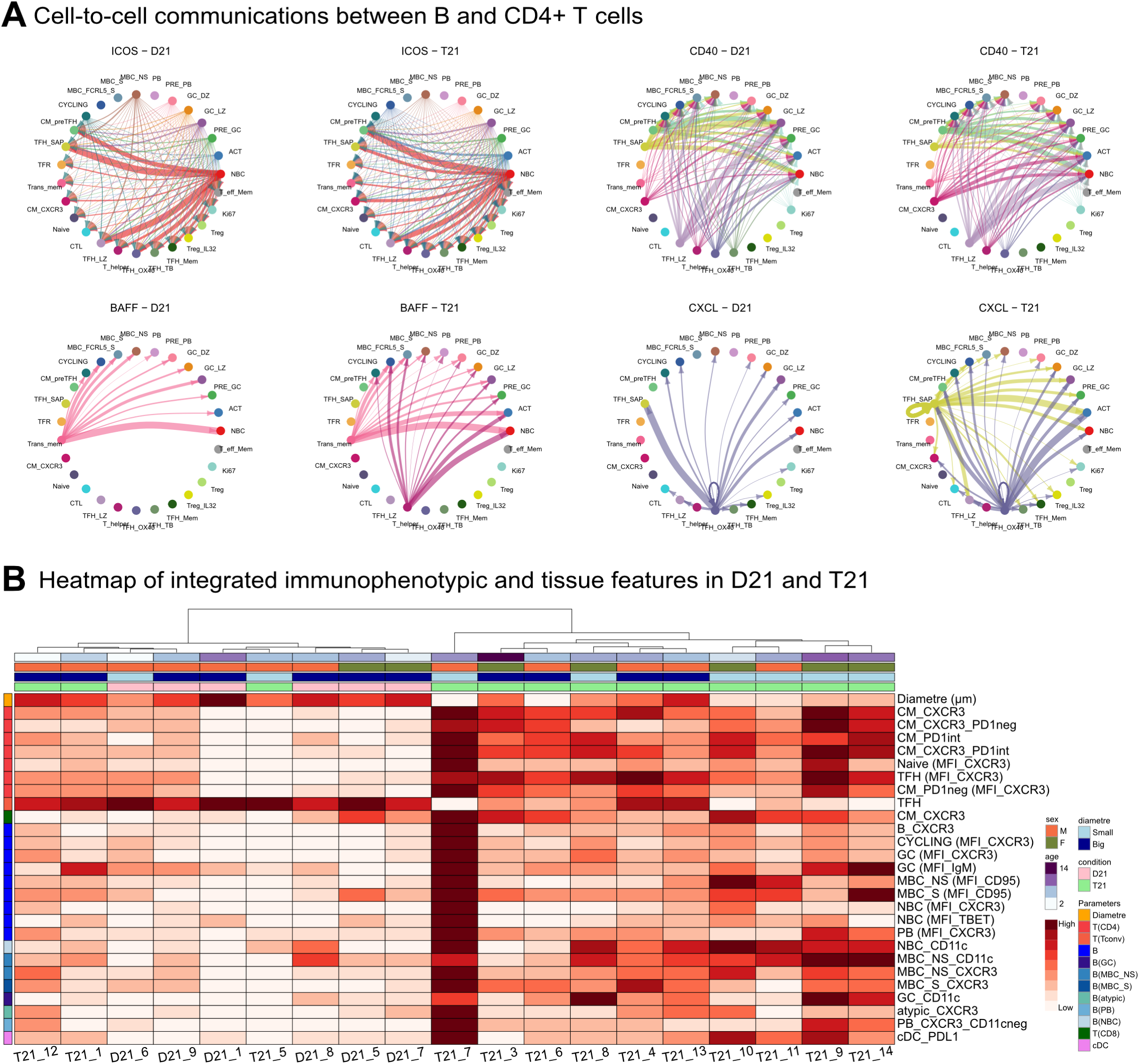
Functional, spatial and integrated characterization of altered T–B interactions in T21 tonsils. **(A)** Predicted cell-cell communication pathways between CD4⁺ T cells and B cells. CellChat analyses depicting ICOS, CD40, BAFF and CXCL signaling networks in D21 and T21 tonsils. **(B)** Integrated heatmap of immunophenotypic and tissue features in D21 and T21 tonsils. Hierarchical clustering integrating follicular measurements, flow cytometric variables and immune cell phenotypes. Samples segregated according to genotypes, ages and follicular size, revealing associations between reduced follicular diameter, increased CXCR3 expression across T- and B-cell populations, and altered memory/helper cell states in T21 tonsils.

**Supplementary Table 1:** Sample allocation across experimental approaches.

|  |  |  |  |  |  |  |  |  |
| --- | --- | --- | --- | --- | --- | --- | --- | --- |
| Sample ID | Diagnosis | Flow |  |  |  | CITE-seq/scTCR/BCR | Diametre measurement | miF |
|  |  | Panel 1 | Panel 2 | Panel 3 | Panel 4 |  |  |  |
| D21_01 | D21 | ✓ | ✓ | ✓ | ✓ |  | ✓ |  |
| D21_03 | D21 | ✓ | ✓ | ✓ | ✓ |  |  |  |
| D21_05 | D21 | ✓ | ✓ | ✓ | ✓ | ✓ | ✓ | ✓ |
| D21_06 | D21 | ✓ | ✓ | ✓ | ✓ |  | ✓ | ✓ |
| D21_07 | D21 | ✓ | ✓ | ✓ | ✓ | ✓ | ✓ | ✓ |
| D21_08 | D21 | ✓ | ✓ | ✓ | ✓ | ✓ | ✓ |  |
| D21_09 | D21 | ✓ | ✓ | ✓ | ✓ |  | ✓ |  |
| D21_10 | D21 |  |  |  | ✓ |  | ✓ |  |
| T21_01 | T21 | ✓ | ✓ | ✓ | ✓ |  | ✓ |  |
| T21_02 | T21 | ✓ | ✓ |  | ✓ |  |  |  |
| T21_03 | T21 | ✓ | ✓ | ✓ | ✓ |  | ✓ | ✓ |
| T21_04 | T21 | ✓ | ✓ | ✓ | ✓ |  | ✓ | ✓ |
| T21_05 | T21 | ✓ | ✓ | ✓ | ✓ |  | ✓ |  |
| T21_06 | T21 | ✓ | ✓ | ✓ | ✓ |  | ✓ | ✓ |
| T21_07 | T21 | ✓ | ✓ | ✓ | ✓ |  | ✓ | ✓ |
| T21_08 | T21 | ✓ | ✓ | ✓ | ✓ | ✓ | ✓ |  |
| T21_09 | T21 | ✓ |  | ✓ | ✓ | ✓ | ✓ |  |
| T21_10 | T21 | ✓ |  | ✓ | ✓ |  | ✓ |  |
| T21_11 | T21 | ✓ |  | ✓ | ✓ |  | ✓ |  |
| T21_12 | T21 | ✓ | ✓ | ✓ | ✓ |  | ✓ |  |
| T21_13 | T21 | ✓ | ✓ | ✓ | ✓ |  | ✓ |  |
| T21_14 | T21 | ✓ | ✓ | ✓ | ✓ | ✓ | ✓ |  |
| T21_15 | T21 |  |  |  | ✓ |  | ✓ |  |
| T21_16 | T21 |  |  |  |  |  | ✓ |  |
| T21_17 | T21 |  |  |  |  |  | ✓ |  |

**Supplementary Table 2:**
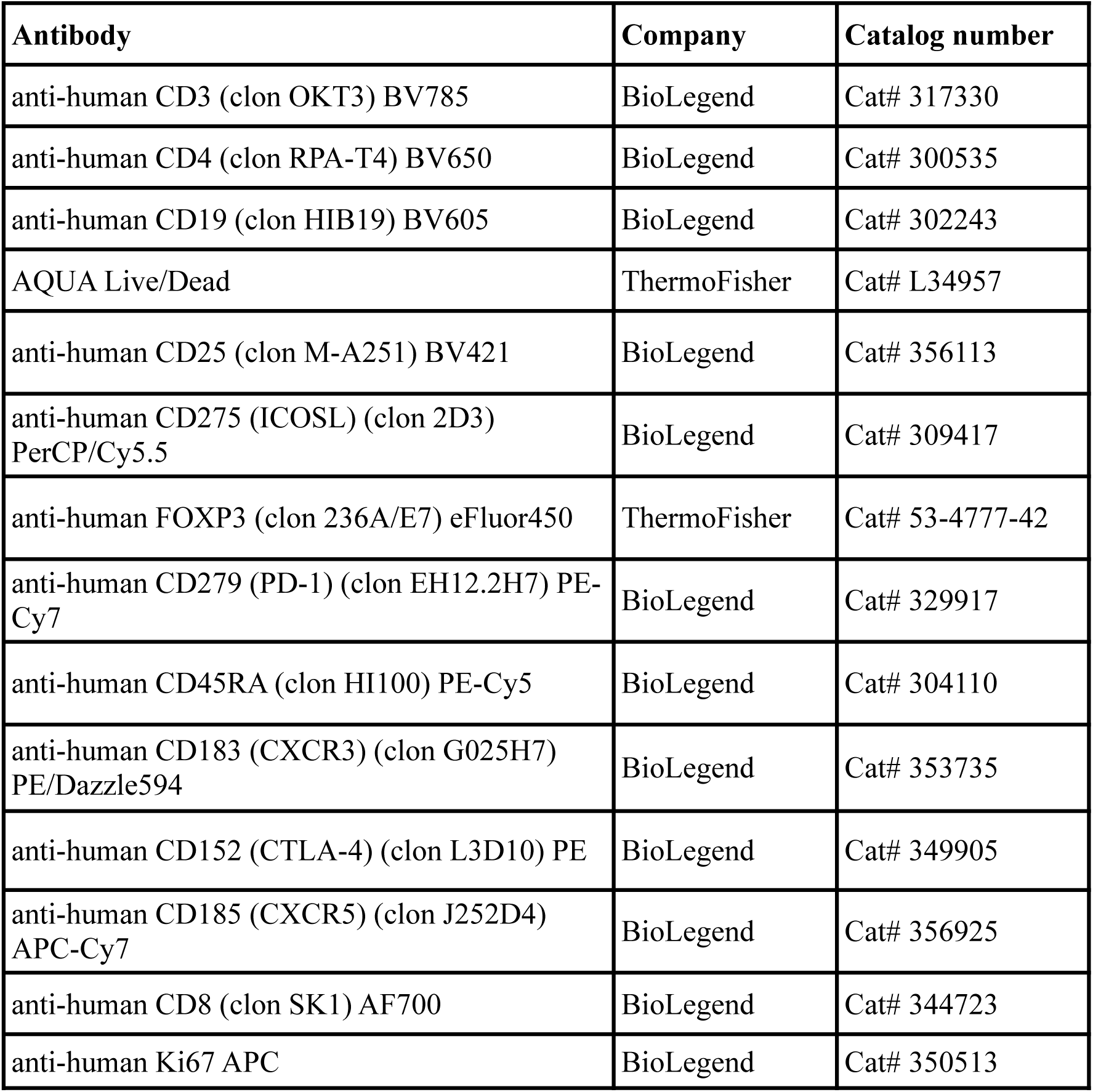
Antibody panel used to characterized T cells by flow cytometry (Panel 1)

| **Antibody** | **Company** | **Catalog number** |
| --- | --- | --- |
| anti-human CD3 (clon OKT3) BV785 | BioLegend | Cat# 317330 |
| anti-human CD4 (clon RPA-T4) BV650 | BioLegend | Cat# 300535 |
| anti-human CD19 (clon HIB19) BV605 | BioLegend | Cat# 302243 |
| AQUA Live/Dead | ThermoFisher | Cat# L34957 |
| anti-human CD25 (clon M-A251) BV421 | BioLegend | Cat# 356113 |
| anti-human CD275 (ICOSL) (clon 2D3) PerCP/Cy5.5 | BioLegend | Cat# 309417 |
| anti-human FOXP3 (clon 236A/E7) eFluor450 | ThermoFisher | Cat# 53-4777-42 |
| anti-human CD279 (PD-1) (clon EH12.2H7) PE-Cy7 | BioLegend | Cat# 329917 |
| anti-human CD45RA (clon HI100) PE-Cy5 | BioLegend | Cat# 304110 |
| anti-human CD183 (CXCR3) (clon G025H7) PE/Dazzle594 | BioLegend | Cat# 353735 |
| anti-human CD152 (CTLA-4) (clon L3D10) PE | BioLegend | Cat# 349905 |
| anti-human CD185 (CXCR5) (clon J252D4) APC-Cy7 | BioLegend | Cat# 356925 |
| anti-human CD8 (clon SK1) AF700 | BioLegend | Cat# 344723 |
| anti-human Ki67 APC | BioLegend | Cat# 350513 |

**Supplementary Table 3:** Antibody panel used to measure cytokine production by intracellular staining (Panel 2)

| **Antibody** | **Company** | **Catalog number** |
| --- | --- | --- |
| anti-human CD3 (clon OKT3) BV785 | BioLegend | Cat# 317330 |
| anti-human IL17A (clon BL168) BV711 | BioLegend | Cat# 512327 |
| anti-human CD4 (clon RPA-T4) BV650 | BioLegend | Cat# 300535 |
| anti-human CD19 (clon HIB19) BV605 | BioLegend | Cat# 302243 |
| AQUA Live/Dead | ThermoFisher | Cat# L34957 |
| anti-human IFNγ (clon 4S.B3) BV421 | BioLegend | Cat# 502531 |
| anti-human CD275 (ICOSL) (clon 2D3)<br>PerCP/Cy5.5 | BioLegend | Cat# 309417 |
| anti-human KLRG1 (clon 13F12F2) Alexa<br>Fluor 488 | ThermoFisher | Cat# 53-9488-42 |
| anti-human CD279 (PD-1) (clon EH12.2H7)<br>PE-Cy7 | BioLegend | Cat# 329917 |
| anti-human CD45RA (clon HI100) PE-Cy5 | BioLegend | Cat# 304110 |
| anti-human GzmB (clon GB11) PE-CF594 | BD Bioscience | Cat# 562462 |
| anti-human IL21 (clon 3A3-N2) PE | BioLegend | Cat# 513003 |
| anti-human CD185 (CXCR5) (clon J252D4)<br>APC-Cy7 | BioLegend | Cat# 356925 |
| anti-human CD8 (clon SK1) AF700 | BioLegend | Cat# 344723 |

**Supplementary Table 4:** Antibody panel used to characterized myeloid immune populations by flow cytometry (Panel 3)

| **Antibody** | **Company** | **Catalog number** |
| --- | --- | --- |
| monoclonal anti-human CD16 (clon 3G8) BV785 | BioLegend | Cat# 302045 |
| monoclonal anti-human CD86 (clon IT2.2) BV711 | BioLegend | Cat# 305439 |
| monoclonal anti-human CD11c (clon Bu15) BV650 | BioLegend | Cat# 337237 |
| monoclonal anti-human HLA-DR (clon L243) BV605 | BioLegend | Cat# 307639 |
| AQUA – Live dead | ThermoFisher | Cat# L34957 |
| monoclonal anti-human FcεRI (clon AER-37) BV421 | BioLegend | Cat# 334623 |
| monoclonal anti-human CD1c (clon L161) PerCP-eFluor710 | ThermoFisher | Cat# 46-0015-42 |
| monoclonal anti-human CD14 (clon 61D3) FITC | ThermoFisher | Cat# 11-0149-42 |
| monoclonal anti-human CD141 (clon M80) PE-Cy7 | BioLegend | Cat# 344109 |
| monoclonal anti-human CD56 (clon 5.1H11) PE-Cy5 | BioLegend | Cat# 362515 |
| monoclonal anti-human CD274 (PD-L1) (clon MIH1) PE-CF594 | BD Bioscience | Cat# 563742 |
| monoclonal anti-human CD304 (clon 12C2) PE | BioLegend | Cat# 354503 |
| monoclonal anti-human CD45 (clon 2D1) APC-Cy7 | BioLegend | Cat# 368515 |
| monoclonal anti-human CD3 (clon OKT3) AF700 | BioLegend | Cat# 317339 |
| monoclonal anti-human CD19 (clon HIB19) AF700 | BioLegend | Cat# 302225 |
| monoclonal anti-human CD370 (CLEC9A) (clon 8F9) APC | BioLegend | Cat# 353805 |

**Supplementary Table 5:**
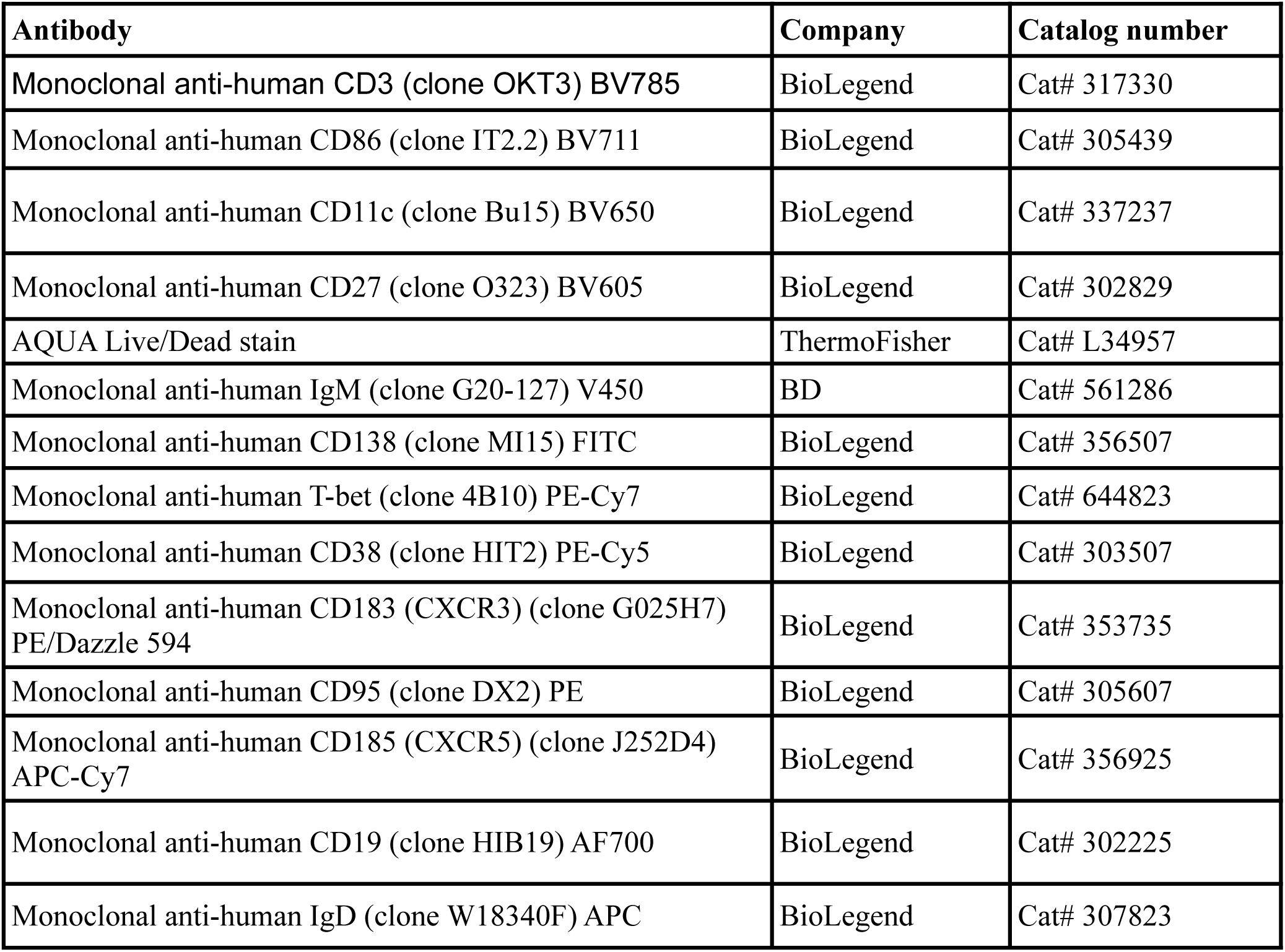
Antibody panel used to characterized B cells by flow cytometry (Panel 4)

| **Antibody** | **Company** | **Catalog number** |
| --- | --- | --- |
| Monoclonal anti-human CD3 (clone OKT3) BV785 | BioLegend | Cat# 317330 |
| Monoclonal anti-human CD86 (clone IT2.2) BV711 | BioLegend | Cat# 305439 |
| Monoclonal anti-human CD11c (clone Bu15) BV650 | BioLegend | Cat# 337237 |
| Monoclonal anti-human CD27 (clone O323) BV605 | BioLegend | Cat# 302829 |
| AQUA Live/Dead stain | ThermoFisher | Cat# L34957 |
| Monoclonal anti-human IgM (clone G20-127) V450 | BD | Cat# 561286 |
| Monoclonal anti-human CD138 (clone MI15) FITC | BioLegend | Cat# 356507 |
| Monoclonal anti-human T-bet (clone 4B10) PE-Cy7 | BioLegend | Cat# 644823 |
| Monoclonal anti-human CD38 (clone HIT2) PE-Cy5 | BioLegend | Cat# 303507 |
| Monoclonal anti-human CD183 (CXCR3) (clone G025H7) PE/Dazzle 594 | BioLegend | Cat# 353735 |
| Monoclonal anti-human CD95 (clone DX2) PE | BioLegend | Cat# 305607 |
| Monoclonal anti-human CD185 (CXCR5) (clone J252D4) APC-Cy7 | BioLegend | Cat# 356925 |
| Monoclonal anti-human CD19 (clone HIB19) AF700 | BioLegend | Cat# 302225 |
| Monoclonal anti-human IgD (clone W18340F) APC | BioLegend | Cat# 307823 |

**Supplementary Table 6:** Antibody panel used to characterized immune cells by multiplex immunofluoerscence (mIF)

| **Antibody** | **Vendor** | **Catalog number** | **Clone** | **pH** | **Dilution** | **Opal** | **Opal dilution** |
| --- | --- | --- | --- | --- | --- | --- | --- |
| HuCD3 | Leica | PA0553 | LN10 | 9 | RTU | 480 | 1:50 |
| HuCD4 | Leica | PA0427 | 4B12 | 9 | RTU | 540 | 1:50 |
| HuBcl6 | Abcam | ab249707 | EPR11410-43 | 9 | 1:250 | 690 | 1:50 |
| HuPD-1 | Abcam | ab52587 | NAT105 | 6 | 1:200 | 650 | 1:150 |
| HuT-bet | CST | 97135S | E4I2K | 6 | 1:200 | 570 | 1:150 |
| HuCXCR3 | Abcam | AB64714 | 49801 | 6 | 1:200 | 520 | 1:50 |
| HuCD21 | Leica | NCL-L-CD21-2G9 | 2G9 | 6 | 1:200 | 620 | 1:50 |
| HuKi67 | EPREDIA (Thermofisher) | RM-9106-S | SP6 | 6 | 1:400 | 780, DIG | 1:25; 1:100 |
| - | Akoya | - | - | - | - | DAPI | 1 |

